# Medial prefrontal dopamine dynamics reflect allocation of selective attention

**DOI:** 10.1101/2024.03.04.583245

**Authors:** Patrick R. Melugin, Suzanne O. Nolan, Evelyn Kandov, Carson F. Ferrara, Zahra Z. Farahbakhsh, Cody A. Siciliano

## Abstract

The mesocortical dopamine system is comprised of midbrain dopamine neurons that predominantly innervate the medial prefrontal cortex (mPFC) and exert a powerful neuromodulatory influence over this region^1,2^. mPFC dopamine activity is thought to be critical for fundamental neurobiological processes including valence coding and decision-making^3,4^. Despite enduring interest in this pathway, the stimuli and conditions that engage mPFC dopamine release have remained enigmatic due to inherent limitations in conventional methods for dopamine monitoring which have prevented real-time *in vivo* observation^5^. Here, using a fluorescent dopamine sensor enabling time-resolved recordings of cortical dopamine activity in freely behaving mice, we reveal the coding properties of this system and demonstrate that mPFC dopamine dynamics conform to a selective attention signal. Contrary to the long-standing theory that mPFC dopamine release preferentially encodes aversive and stressful events^6–8^, we observed robust dopamine responses to both appetitive and aversive stimuli which dissipated with increasing familiarity irrespective of stimulus intensity. We found that mPFC dopamine does not evolve as a function of learning but displays striking temporal precedence with second-to-second changes in behavioral engagement, suggesting a role in allocation of attentional resources. Systematic manipulation of attentional demand revealed that quieting of mPFC dopamine signals the allocation of attentional resources towards an expected event which, upon detection triggers a sharp dopamine transient marking the transition from decision-making to action. The proposed role of mPFC dopamine as a selective attention signal is the first model based on direct observation of time-resolved dopamine dynamics and reconciles decades of competing theories.

## Main Text

The mesocortical dopamine system, comprised of dopamine releasing terminals in prefrontal cortex arising from the midbrain, was described just a few years after the discovery of central dopamine in the canonical mesolimbic and nigrostriatal dopamine circuits^9–11^. Similar to the subcortical dopamine systems, mesocortical dopamine has been the subject of intensive research efforts since its discovery; however, insight into the neurobiological functions of cortical dopamine have remained notoriously elusive. The relative paucity of studies investigating dopamine in the medial prefrontal cortex (mPFC) results from the neurochemical heterogeneity of this region, where dopamine and norepinephrine are released from neighboring boutons^2,12,13^. Conventionally, real-time observation of dopamine release in intact tissue is achieved exclusively via electrochemical methodologies, such as fast-scan cyclic voltammetry, which cannot distinguish dopamine from norepinephrine; this inability to distinguish catecholamines has long precluded real-time electrochemical monitoring of mPFC dopamine dynamics^5^. Accordingly, only a handful of studies have obtained time-resolved measurements of mPFC dopamine release *in vivo*. These studies were performed in anesthetized animals and for unambiguous interpretation required that dopamine release was evoked via stimulation of dopamine soma in the midbrain^14,15^ or required extensive *post-hoc* control experiments^16^. Recent advances in fluorescent biosensors permit selective dopamine monitoring with millisecond resolution^17,18^, potentially circumventing roadblocks with electrochemical approaches. Here, we leveraged fluorescent dopamine sensing to directly interrogate the functional properties of the system during ongoing behavior.

While the importance of the mesocortical dopamine system in adaptive behaviors and neuropsychiatric disease states is undisputed, there is little consensus as to the precise coding properties of this system. The longest standing theory of mesocortical dopamine system function is that, in contrast to the mesolimbic dopamine system, this circuit is selectively responsive to stressful and aversive stimuli^6,8^. Support for this theory comes largely from electrophysiological measures of somatic action potential activity in midbrain dopamine neurons projecting to the mPFC which display tail pinch-evoked activity *in vivo*^7^ and increased synaptic strength *ex vivo* following exposure to noxious stimuli^19,20^. Further, tissue content of dopamine metabolites are augmented following exposure to aversive stimuli^21–25^. In contrast, assessments of extracellular dopamine concentrations in the mPFC measured via microdialysis, which allows selective dopamine quantification but low temporal resolution on the order of tens of minutes, have revealed increased dopamine activity after exposure to stimuli with both positive and negative valence^26–29^, and competing theories posit that mPFC dopamine has more complex roles in higher order cognition and attentional processes^30–32^. Due to the fact that previous studies did not have sufficient temporal resolution to resolve the precise behavioral events associated with dopamine elevations, little progress has been made in unifying these seemingly disparate views of mPFC dopamine’s function.

### Stimulus-evoked mPFC Dopamine Transients do not Differentiate Valence or Intensity

Given the challenges of implementing previous approaches in cortex, we first sought to directly verify whether a fluorescent biosensor strategy could provide sufficient sensitivity and selectivity for unambiguous dopamine monitoring in the mPFC of awake, freely behaving mice. Fluorescent dopamine biosensors are based on endogenous dopamine receptors, with various mutations introduced to couple dopamine binding to fluorophore conformation, and an expanding range of variants are currently available^17,18^. We selected the dLight family of fluorescent dopamine sensors which are D1 receptor-based, given the relatively low affinity of endogenous D1 receptors for norepinephrine. Given that the peak concentrations of extracellular dopamine in mPFC during coordinated release events is unknown, dLight1.2 was an attractive choice among the available variants as it displays wide dynamic range while retaining high sensitivity, allowing for scaled responses from the low nanomolar to mid-micromolar range^17^. In mPFC acute *ex vivo* slices expressing dLight1.2 (**Extended Data Fig. 1a**), we reproduce the results of Patriarchi and colleagues^17^ measured in cultured cells, demonstrating that dLight1.2 responds to dopamine in the low nanomolar range and scales in fluorescent intensity up to at least 100 µM when excited with blue (490nm center wavelength) light. Further, we find that UV spectrum excitation (405nm center wavelength) is isosbestic and displays minimal changes in fluorescence intensity over the same dopamine concentration range (**Extended Data Fig. 1b,c**). Critically, dLight1.2 exhibits high selectivity for dopamine over norepinephrine, as neither 490 nor 405nm excited fluorescence displayed appreciable changes in fluorescence intensity in response to norepinephrine at concentrations below 100 µM (**Extended Data Fig. 1**).

To leverage this sensor for the monitoring of *in vivo* dopamine dynamics, we injected a viral vector encoding dLight1.2 into the mPFC and implanted a chronic indwelling fiber optic cannula (**Fig. 1a; Extended Data** Fig. 2) in order to perform fiber photometry in freely moving mice (**Fig. 1b**). Given that the standing theories of mPFC dopamine function are largely based on recordings in anesthetized animals, our initial investigations focused on discrete, unconditioned stimuli to facilitate comparison. Consistent with claims that mPFC dopamine preferentially responds to aversive stimuli, we observed a robust dopamine response to tail pinch in non-anesthetized animals, which was qualitatively similar to previously reported dopamine-verified electrochemical recordings in anesthetized animals (**Fig. 1c**) (c.f. ^16^). Tail pinch-evoked transients were markedly reduced following administration of a D1/dLight receptor antagonist, confirming that fluorescent signals were dependent on dopamine-dLight binding **(Fig 1d; Extended Data Fig. 3a)**.

**Figure 1.**
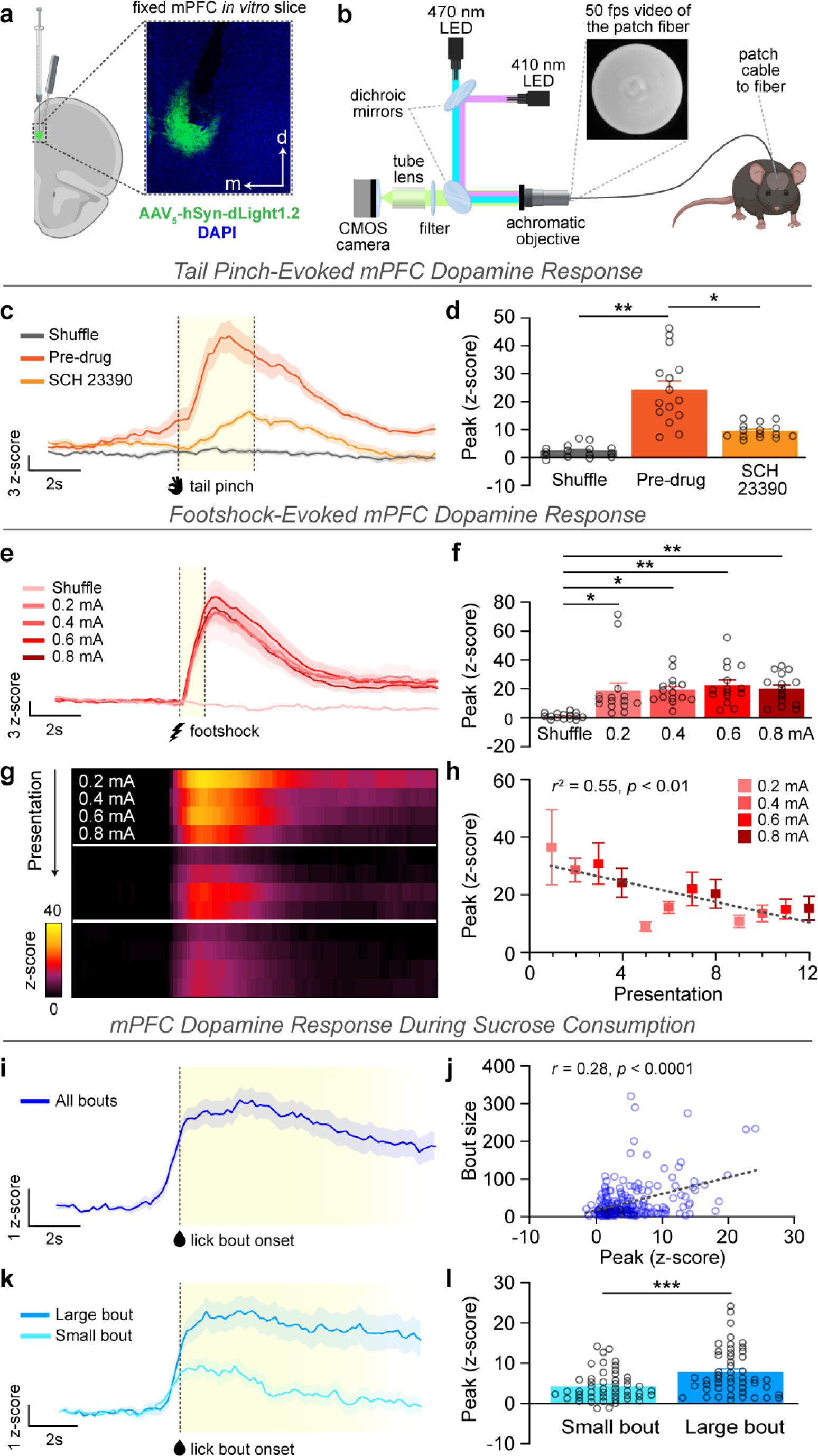
Stimulus-evoked mPFC dopamine transients do not differentiate stimulus valence. **(a)** Representative histological image showing dLight1.2 expression and fiber optic implant placement in the medial prefrontal cortex (mPFC). **(b)** Schematic of fiber photometry setup used to record fluctuations in dLight fluorescence in the mPFC of behaving mice. **(c)** dLight fluorescence intensity traces indicating mPFC dopamine activity over time aligned to tail pinch onset under baseline conditions (pre-drug) and following blockade of dLight/D1 receptors via SCH 23390 (1 mg/kg i.p.). A time-shuffled alignment, where signal was aligned to a pseudorandomly selected time during the interstimulus period, was used to determine signal observed by chance. Vertical lines indicate tail pinch onset and offset (n = 30 trials, sampled from 3 subjects). **(d)** The magnitude of the tail pinch-evoked response was greater than the shuffled time alignment and was attenuated by systemic delivery of the dLight antagonist (nested ANOVA, *F*_(2,6)_ = 15.86, *p* = 0.0040; Tukey’s test, Shuffle vs. Pre-drug, *p* = 0.0036; Shuffle vs. SCH 23390, *p* = 0.2414; Pre-drug vs. SCH 23390, *p* = 0.0234). **(e)** Dopamine responses to unpredictable footshocks presented in a series of ascending intensity (0.2 – 0.8 mA in 2 mA steps, 1s duration, delivered under a variable-time 30s schedule) with shuffled time alignment comparison. Vertical lines indicate footshock onset and offset. This series was repeated in triplicate (n = 60 trials, sampled from 5 subjects). **(f)** Footshock-evoked responses were greater than the shuffled time alignment but did not differ as a function of amperage (nested ANOVA, *F*_(4,20)_ = 6.028, *p* = 0.0024; Tukey’s test, Shuffle vs. 0.2mA, *p* = 0.0158; Shuffle vs. 0.4mA, *p* = 0.0125; Shuffle vs. 0.6mA, *p* = 0.0027; Shuffle vs. 0.8mA, *p* = 0.0092; *p* > 0.05 for all other comparisons). **(g)** Heatmap displaying dopamine activity (z-axis) averaged across animals for each of the 12 footshock presentations (y-axis), aligned around footshock onset (x-axis). **(h)** Footshock-triggered dopamine transients decreased as a function of presentation order regardless of amperage (simple linear regression, *r*^2^ = 0.5528, *F*_(1,10)_ = 12.36, *p* = 0.0056). **(i)** Dopamine activity during consumption of a sucrose solution (10% w/v), aligned around lick bout onset (n = 200 lick bouts, sampled from 6 subjects). **(j)** The number of licks in each bout was positively correlated with the magnitude of the dopamine response (Spearman’s correlation, *r* = 0.2799, *p* < 0.0001). **(k)** Dopamine traces associated with large (upper quartile) and small (lower quartile) bout sizes. **(l)** Bouts with a higher number of licks produced a larger dopamine response (Mann-Whitney U test, *U* = 776, *p* = 0.0010). Data represented as mean ± S.E.M. * *p* < 0.05; ** *p* < 0.01; *** *p* < 0.001.

Having replicated the results of the previous study that directly measured real-time stimulus-evoked mPFC dopamine release in anesthetized animals^16^, we next sought to determine whether the proposed theories are consistent across aversive stimulus modality and intensity. Mice were tested in operant chambers where they were exposed to a series of unsignaled footshocks of increasing amperage (0.2 - 0.8 mA, ascending, series repeated in triplicate) delivered on a variable time schedule. Similar to tail pinch, footshock evoked a large dopamine response (**Fig. 1e**). However, the magnitude of the dopamine response did not differ as a function of footshock amperage, though there was considerable across-trial variability (**Fig. 1f; Extended Data Fig. 3b**). Additional analysis revealed that, contrary to our hypothesis that the dopamine response would scale with the amperage of the shock, variance across trials was instead largely explained by the number of times the subject experienced the stimulus (**Fig. 1g**). Indeed, there was a linear decrease in the magnitude of the dopamine response across subsequent footshocks irrespective of intensity (**Fig 1h; Extended Data Fig. 3c**). These results suggest that mPFC dopamine may also be encoding features of stimulus familiarity.

While the previous results thus far corroborate mPFC dopamine’s putative involvement in aversive processing, falsifiability of this theory has not been possible due to the lack of established approaches for delivering appetitive stimuli in anesthetized animals. To empirically test if mPFC dopamine is preferentially responsive to aversive over appetitive stimuli, mice were given access to a sipper tube containing a sucrose solution, and dopamine activity was aligned to the initiation of lick bouts. Contrary to standing theory, mPFC dopamine transients were observed around the initiation of sucrose lick bouts (**Fig. 1i**). Sucrose lick bout associated mPFC dopamine transients were reliably observed across many bouts and scaled in magnitude with bout size and duration (**Fig. 1j-l; Extended Data Fig. 3d-f**). Interestingly, the rise of the mPFC dopamine signaling was evident just prior to first lick contact in a bout. Further analysis of continuous timeseries data, without aligning around specific behavioral events, revealed striking covariance between mPFC dopamine activity and lick rate on a sub-second timescale, again with dopamine transients tending to slightly precede bout initiation as well as within-bout variance in lick bursts (**Extended Data Fig. 4; Supplemental Video 1**). These results demonstrate that mPFC dopamine does not distinguish stimuli based on valence and therefore falsifies a leading theory of mPFC dopamine functionality.

The data above clearly does not support valence coding as an explanatory construct for mPFC dopamine functionality, but also does not clearly indicate a plausible alternative. We speculated that these data may indicate a role in novelty processing, as evidenced by the presentation-dependent decrease in mPFC dopamine response to footshock, and behavioral engagement as evidenced by the second-to-second co-variance between mPFC dopamine activity and lick rate. We next sought to directly test whether mPFC dopamine activity tracks novelty/habituation processes in the absence of discrete stimuli. Experimentally naïve mice were placed into an operant chamber with which they had no prior experience and allowed to explore the context, without any experimental manipulations, during 2 consecutive daily sessions (**Fig. 2a**). Spontaneous dopamine transients were observed across both the novel (day 1) and familiar (day 2) sessions (**Fig. 2b**). Strikingly, the occurrence of spontaneous events was exceedingly infrequent (**Fig 2c**), occurring orders of magnitude less frequently than has been reported in striatal dopamine circuits^33,34^. Consistent with our hypothesis regarding novelty processing, the frequency of spontaneous events was higher in the novel vs. familiar context (**Fig. 2c**,**d**).

**Figure 2.**
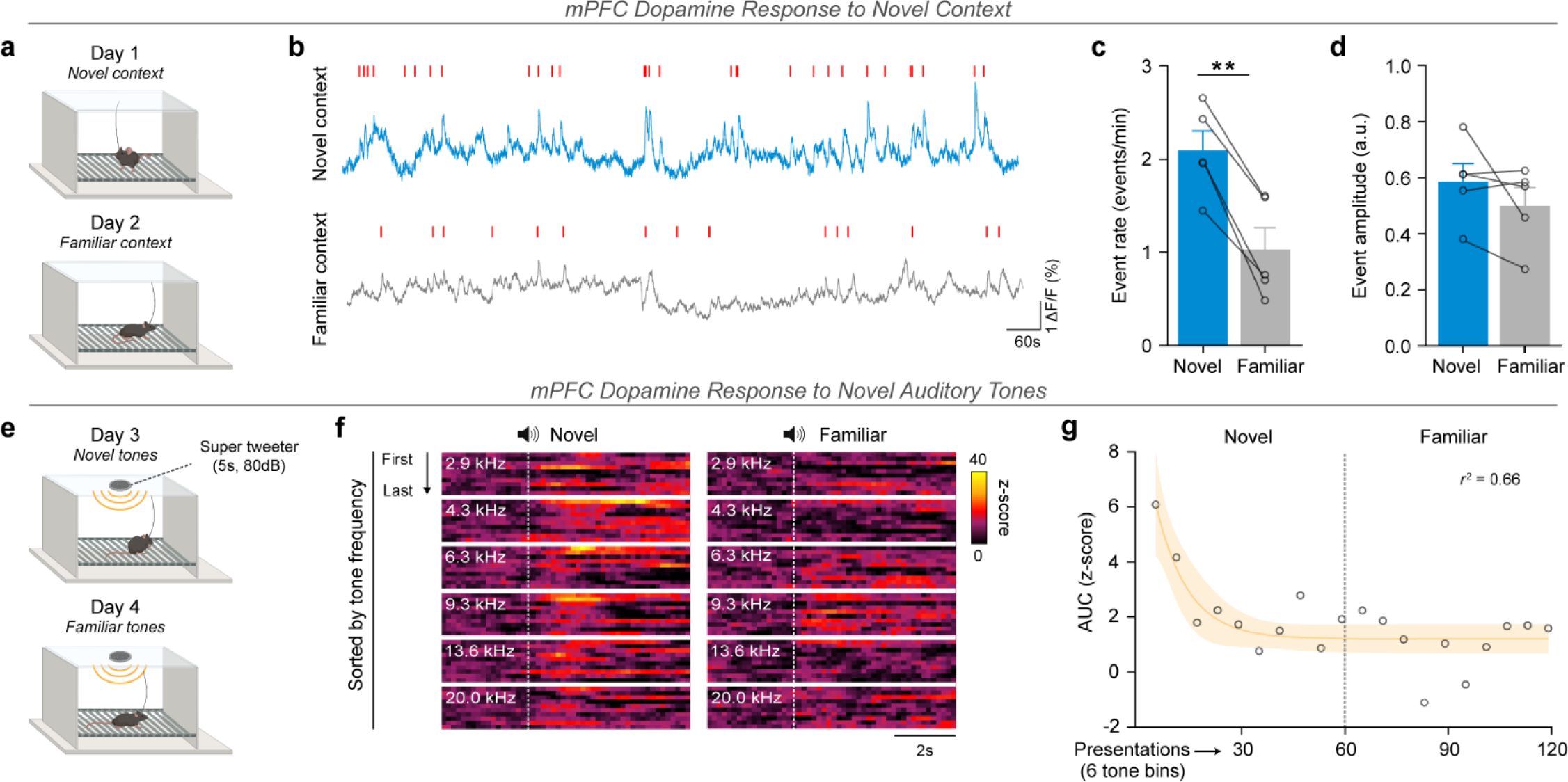
Novel contexts and stimuli engage mPFC dopamine transients which dissipate throughout habituation. **(a)** Schematic of experimental design. Experimentally naïve mice were placed in an unfamiliar operant box devoid of manipulanda for 20-minute sessions, repeated across 2 consecutive days. **(b)** Full-session traces of mPFC dopamine activity from a single animal for both sessions. Red marks correspond to events meeting detection threshold. **(c)** There was a lower frequency of events in the novel context on the second day of exposure compared to the first day (paired samples t-test, *t*_(4)_ = 7.424, *p* = 0.0018), **(d)** but no difference in event amplitude (paired samples t-test, *t*_(4)_ = 1.354, *p* = 0.2472). **(e)** Following habituation to the operant box, animals underwent 2 sessions across consecutive days wherein they were exposed to 6 pure auditory tones presented on a variable-time-15s schedule (2.9 kHz – 20.0 kHz in two-thirds-octave steps, 5s duration, 80dB). The 6 tones were presented in a pseudorandom block design (10 blocks per session, 60 presentations per session; total n = 300 trials recorded per session, sampled from 5 subjects). **(f)** Heatmaps displaying dopamine activity (z-axis) averaged across animals for each tone presentation (y-axis), aligned around tone onset (x-axis). **(g)** Best non-linear fit (*r*^2^ = 0.66) shown with a 95% confidence band demonstrating a decay in dopamine response across presentations. Individual points represent the average response for each block of 6 tone presentations. The span of the curve (upper minus lower plateau) was elevated relative to zero (one sample t-test, H_0_ = 0, *t*_(4)_ = 5.418, *p* = 0.0056). Data represented as mean ± S.E.M. ** *p* < 0.01.

Next, we sought to determine if mPFC dopamine tracks stimulus familiarity for discrete stimuli without clear positive or negative valence. Following novel context exposure, animals underwent 2 additional daily sessions in the now familiar context where they were exposed to pure auditory tones comprising 6 distinct frequencies (2.9 – 20 kHz), presented in a pseudorandom block design (10 blocks of 6 tones per session, random order within block) (**Fig. 2e**). To first test for frequency tuning, we compared dopamine responses to individual tone frequencies collapsed across sessions. We observed a tone-evoked moderate increase in mPFC dopamine activity which did not vary as a function of frequency (**Extended Data Fig. 5**). Splitting into novel and familiar sessions revealed an apparent reduction in tone responsiveness across all frequencies tested (**Fig. 2f**). Analysis of tone-evoked responses by presentation order, irrespective of frequency, revealed an exponential decline in activity occurring throughout the first few tone presentations followed by a plateau near zero throughout the remaining presentations across both sessions (**Fig. 2g**). These data demonstrate that mPFC dopamine is evoked by contextual and discrete stimuli independent of valence or intensity and that these responses decline as a function of stimulus familiarity.

### mPFC Dopamine Dynamics Do Not Evolve Across Reinforcement Learning

While there is considerable evidence that midbrain dopamine neurons innervating the mPFC are anatomically and physiologically distinct from those innervating the ventral striatum^3,35^, the results above raise the question as to the extent to which these systems are in fact functionally distinct in terms of dopamine release patterns. Indeed, the mesolimbic dopamine system is engaged by stimuli with both positive and negative valence, and is modulated by stimulus familiarity^36,37^, largely mirroring the results above. Given that a defining feature of the mesolimbic dopamine system is learning-induced plasticity, particularly regarding learning of stimulus-reinforcer contingencies^36–40^, we next sought to determine whether mPFC dopamine dynamics evolve over the course of reinforcement learning.

Animals were trained in 2 phases, first to acquire an operant reinforced by presentation of sucrose, and next to acquire a discriminated operant whereby an antecedent discriminative stimulus (S^D^) signals when the contingency is in effect. These phases were used to comprehensively test for learning-induced shifts in mPFC dopamine dynamics as a function of basic reinforcement learning, where an action is associated with a positive outcome, and complex learning, where a neutral stimulus acquires value due to its predictive relationship with a primary reinforcer. During the first phase, deemed continuous reinforcement, animals were trained to respond on the active side, denoted by an illuminated cue light above the operandum, which resulted in extension of a sipper containing sucrose solution (**Extended Data Fig. 6a**). Across sessions, mice rapidly learned the contingency as evidenced by increased responding on the active side and lower latency to initiate a lick bout following an active response (**Extended Data** Fig. 6b,c). Further, there was no evidence of familiarity- or satiety-induced changes in response to the primary reinforcer as the number of licks per sucrose access period remained stable (**Extended Data Fig. 6d**). Aligning mPFC dopamine activity around task events (**Extended Data Fig. 6e,f**) revealed no change dopamine activity at the time of reinforced lever press during early and late learning (**Extended Data Fig. 6g**); however, the magnitude of the dopamine response during sucrose consumption was reduced after learning (**Extended Data Fig. 6h**). Consistent with results during open access sucrose consumption (**Fig. 1**), we again observed an initial rise in mPFC dopamine which preceded the first lick contact in the bout, which remained unchanged across learning (**Extended Data Fig. 6**). Analysis of individual animals across all sucrose reinforcers earned throughout the task revealed that reduction was attributable to a gradual, linear decline in mPFC dopamine response magnitude as a function of the number of sucrose access periods but did not differ between fast and slow learners (**Extended Data Fig. 7**). These data show that stimulus familiarity-related changes are again observable across reinforcer presentations but do not provide evidence of learning-related changes across acquisition of basic positive reinforcement.

Next, animals were trained on a discriminated operant reinforcement task. Animals were tested as a continuation of the prior task, without altering the experimental context. In this phase, the only change to the procedure was that the cue light above the active operandum was presented on a variable time schedule and served as a S^D^. Accordingly, only responses made on the active side in the presence of the S^D^ were reinforced by extension of the sucrose sipper. A response on the active side in the absence of the S^D^ (S^Δ^ period) resulted in a 30 second timeout period where no responses were reinforced (**Fig. 3a**). Mice demonstrated a clear divergence over sessions in the number of reinforcers earned relative to timeout periods triggered **(Fig. 3b**).

**Figure 3.**
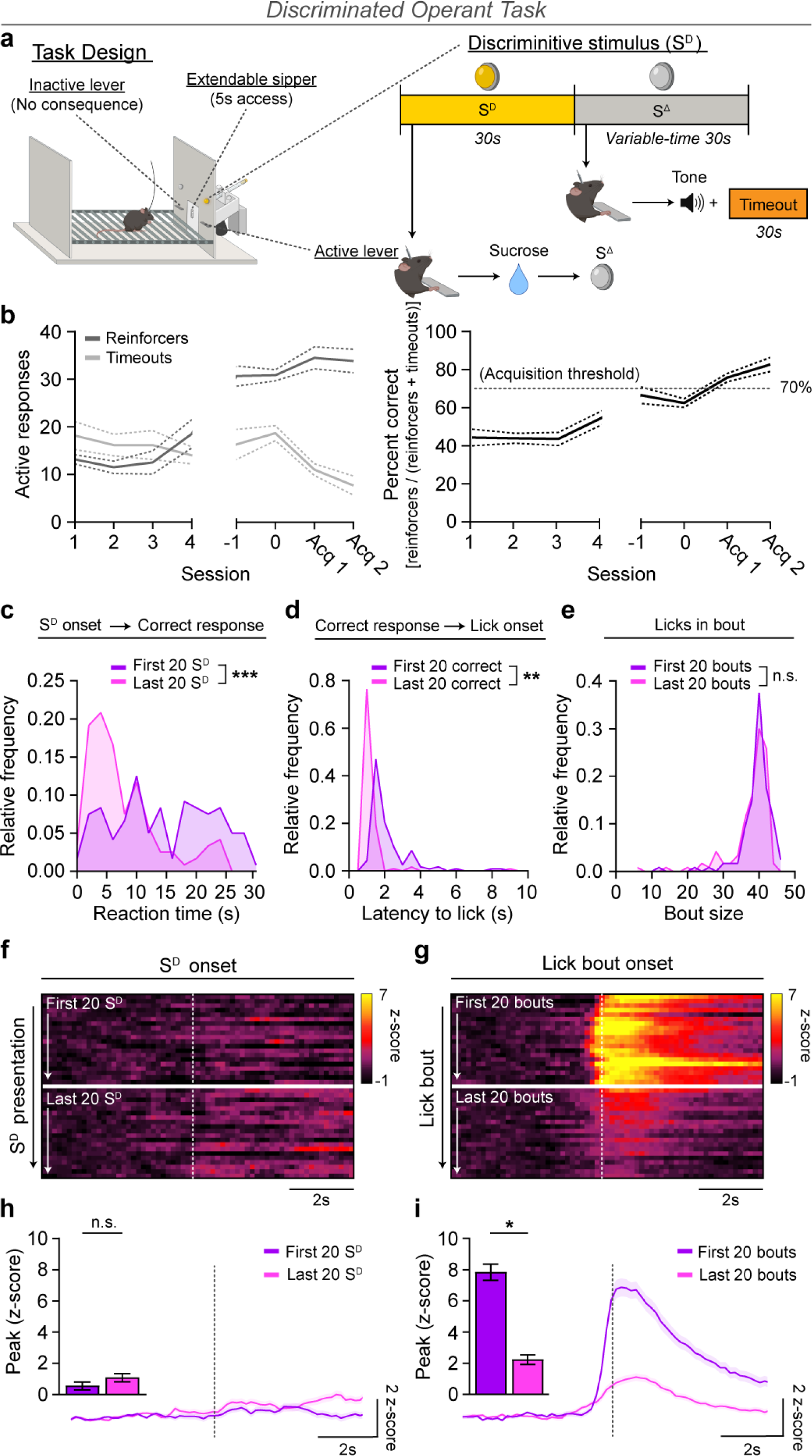
mPFC dopamine dynamics do not evolve over acquisition of complex contingency learning. **(a)** Schematic of discriminated operant learning task. A response on the active operandum in the presence of the discriminative stimulus (S^D^), deemed a correct response, was reinforced by extension of a sipper tube containing 10% sucrose (w/v) which remained accessible for a 5s period commencing with first lick contact. A response on the active operandum in the absence of the S^D^ (S^Δ^ period) triggered a 30s timeout period signaled by the presentation of an auditory tone. **(b)** *Left*: Behavior data demonstrating a divergence in correct responses and timeout responses across sessions. *Right*: Animals were tested until performance reached ≥ 70% correct responses [reinforcers earned/(reinforcers earned + timeouts initiated)] for 2 consecutive sessions, denoted as acquisition day 1 (Acq 1) and day 2 (Acq 2), while attaining a minimum of 20 correct responses in each session. **(c-e)** Comparison of behavioral measures during the first 20 correct trials during the pre-acquisition period and the first 20 correct trials on the last acquisition day. **(c)** By the final session, subjects displayed a faster reaction time to correctly respond following S^D^ onset (nested ANOVA, *F*_(1,10)_ = 28.09, *p* = 0.0003) **(d)** and lower latencies to initiate a lick bout following a correct response (nested ANOVA, *F*_(1,10)_ = 11.12, *p* = 0.0076) **(e)** while exhibiting no difference in the number of licks per bout (nested ANOVA, *F*_(1,10)_ = 0.7874, *p* = 0.3957). **(f)** Heatmap displaying dopamine activity (z-axis) surrounding S^D^ onset (x-axis) averaged across animals for each of the first 20 S^D^ presentations (y-axis) during the pre-acquisition period and the last acquisition day. **(g)** Heatmap displaying averaged dopamine activity (z-axis) surrounding lick bout onset (x-axis) averaged across animals for each of the first 20 lick bouts (y-axis) during the pre-acquisition period and the last acquisition day. **(h)** Averaged dopamine traces (n = 120 events per learning epoch, sampled from 6 subjects). Vertical line indicates S^D^ onset. *Inset:* There was no change in dopamine response to the S^D^ after learning (nested ANOVA, *F*_(1,10)_ = 1.220, *p* = 0.2952). **(g)** Averaged dopamine traces (n = 120 events per learning epoch, sampled from 6 subjects). Vertical line indicates lick bout onset. *Inset:* The magnitude of the dopamine response following lick bout onset decreased after learning (nested ANOVA, *F*_(1,10)_ = 9.057, *p* = 0.0131). Data represented as mean ± S.E.M. * *p* < 0.05; ** *p* < 0.01; *** *p* < 0.001.

Further, over the course of learning mice displayed markedly faster correct responses following presentation of the S^D^ (**Fig. 3c**), exhibited lower latencies to initiate lick bouts following a correct response (**Fig. 3d**), but no change in the number of licks in a bout (**Fig. 3e**). Critically, we did not observe any transfer of the dopamine response at the time of reinforcer receipt to the antecedent S^D^ as would be expected for a learning signal (**Fig. 3f,h**), despite clear behavioral evidence that the previously neutral S^D^ had acquired value (**Fig. 3 b-d**). As expected based on the consistent familiarity-related effects on mPFC dopamine responses outlined above, there was a reduction in the magnitude of dopamine response during sucrose consumption across initial versus post-acquisition trials (**Fig. 3g,i**). Together, these data do not support learning-dependent alterations in mPFC dopamine dynamics, in stark contrast to the mesolimbic system.

### mPFC Dopamine Dynamics Reflect Internal States in the Absence of Discrete Stimuli

Throughout the experiments detailed above, there appears to be two highly consistent findings regarding mPFC dopamine dynamics: 1) familiarity-related decreases in stimulus-evoked activity, and 2) ramping activity prior to the initiation of licking behavior. Novelty processing alone is not sufficient to explain these dynamics, given that the signal can precede the event (**Fig. 1**) and responses do not fully dissipate even after many exposures (**Extended Data Fig. 6, Fig. 3**). In search of a unifying explanatory construct, we next explored whether the observed mPFC dopamine activity prior to sucrose consumption reflects a dissociable component from the apparent response to sucrose itself. To accomplish this, mice that were trained to respond for sucrose access were tested under a variable delay reinforcement contingency, wherein a delay period (0, 2, or 5 seconds) was probabilistically introduced between a correct response and the resulting extension of the sipper tube (**Fig. 4a**). We found that mPFC dopamine activity begins to ramp prior to reinforcer receipt, and that this ramping activity scales as the delay period increases (**Fig. 4b**), demonstrating that this signal is related to an internal state rather than an action *per se*. Further, with longer delays between action (lever press) and outcome (sipper extension), and thus increased dopamine activity prior to sucrose receipt, there was a commensurate decrease in the dopamine response occurring during the sucrose consumption period (**Fig. 4c**). Thus, while the aggregate dopamine response did not differ between trial types (**Fig. 4d**), the distribution of the response during pre- and post-stimulus periods varied as a function of the delay period (**Fig. 4e**).

**Figure 4.**
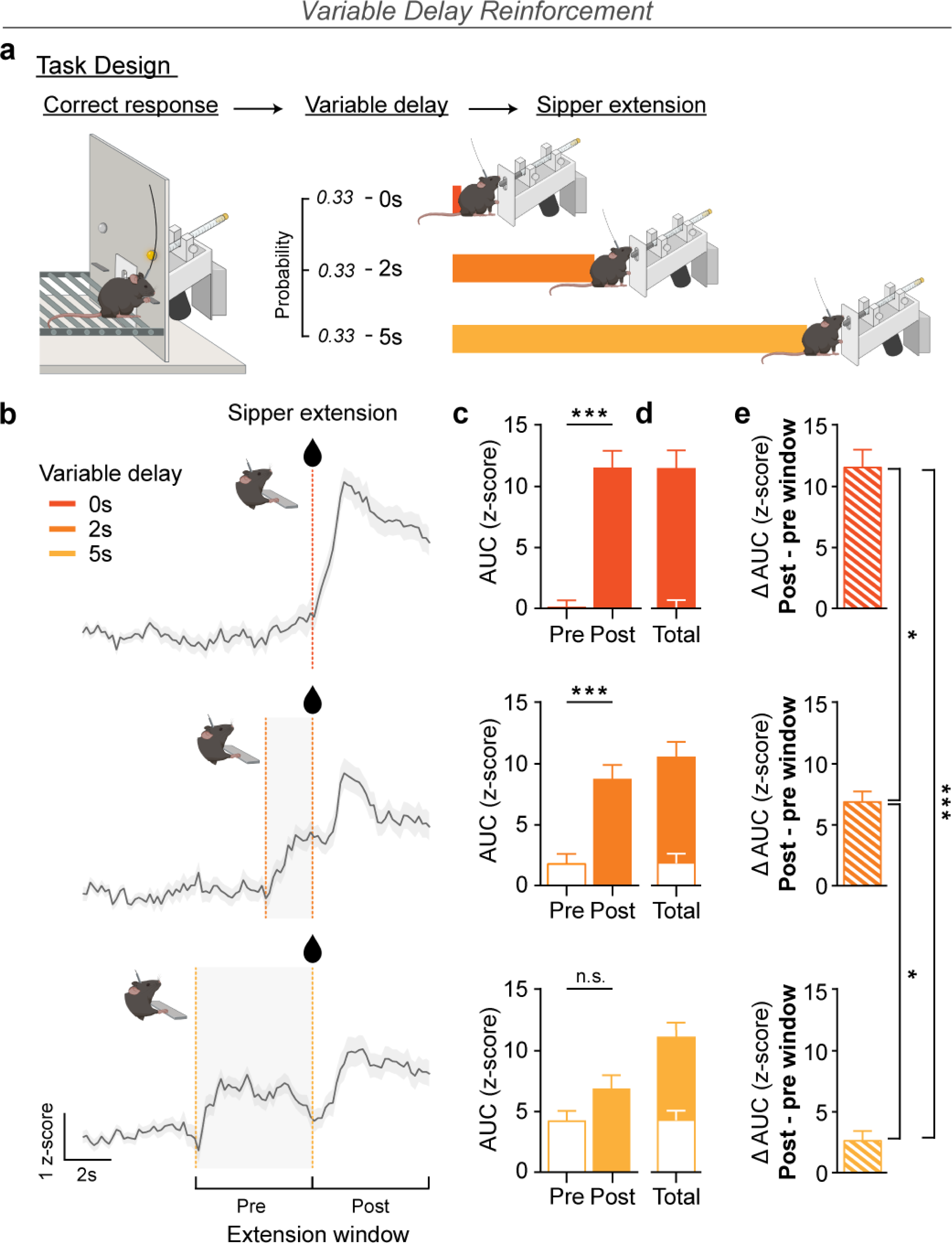
mPFC dopamine activity is engaged during anticipation of delayed reward. **(a)** Schematic of variable delay reinforcement task wherein a delay to the sipper extension (0s, 2s, 5s) was probabilistically introduced following a correct response (pseudorandom order). **(b)** Dopamine activity aligned around the extension of the sipper tube. For no-delay trials (0s), the vertical line indicates both the correct response and the resulting sipper extension. For delay trials (2s, 5s), vertical lines (left to right) indicate the timing of the correct response and the sipper extension following the corresponding delay period. **(c)** Comparisons of dopamine activity during a 5s pre- and 5s post-extension window for each trial type (two-way repeated measures ANOVA, epoch, *F*_(1,199)_ = 112.1, *p* < 0.0001; trial type, *F*_(2,199)_ = 0.0691, *p* = 0.9333; epoch × trial type, *F*_(2,199)_ = 16.07, *p* < 0.0001). Dopamine activity was greater during the post-extension window on no-delay (n = 77, sampled from 5 subjects) and 2s-delay (n = 61) trials but not 5s-delay (n = 64) trials (planned Šídák’s test, 0s delay pre vs post, *p* < 0.0001; 2s delay pre vs post, *p* < 0.0001; 5s delay pre vs post, *p* = 0.0803). **(d)** There was no difference in the aggregate (post + pre-extension window) dopamine activity across trial types (one-way ANOVA, *F*_(2,199)_ = 0.04792, *p* = 0.9532). **(e)** Comparison of the peristimulus difference in dopamine activity (post-minus pre-extension window) across trial types (one-way ANOVA, *F*_(2,199)_ = 15.91, *p* < 0.0001). The disparity in activity between post- and pre-extension windows varied as a function of the size of the delay period (Tukey’s test, 0s vs. 2s, *p* = 0.0112; 0s vs. 5s, *p* < 0.0001; 2s vs. 5s, *p* = 0.0325). Data represented as mean ± S.E.M. * *p* < 0.05; *** *p* < 0.001.

The results of the delayed reinforcement experiment indicate that mPFC dopamine responses before and during reinforcer receipt can be modulated by expectation, even when the properties of the reinforcer itself are invariant. This raises the intriguing possibility that the signals observed during sucrose consumption throughout the experiments above are not causally tied to sucrose itself, and instead may reflect a co-occurring behavioral or internal process. To test this possibility, animals next underwent a single conditioned reinforcement session wherein the task parameters were unchanged, but responding was reinforced by extension of a dry/empty sipper. We then compared the dopamine response surrounding lick bout onset for the first 5 extensions following a correct response, during which mice reliably licked in the absence of sucrose (**Extended Data Fig. 8**). Despite the absence of the primary reinforcer, or any discrete stimulus for that matter, there was a pronounced increase in mPFC dopamine activity beginning just prior to the first lick and continuing throughout the duration of the bout (**Fig. 5a**), mirroring activity observed during licking for sucrose (**Fig. 1, 3; Extended Data Fig. 6**). Consistent with our earlier observations (**Fig. 1; Extended Data Fig. 4**), the magnitude of the dopamine response scaled markedly with the ongoing rate of licking (**Fig. 5b,c**).

**Figure 5.**
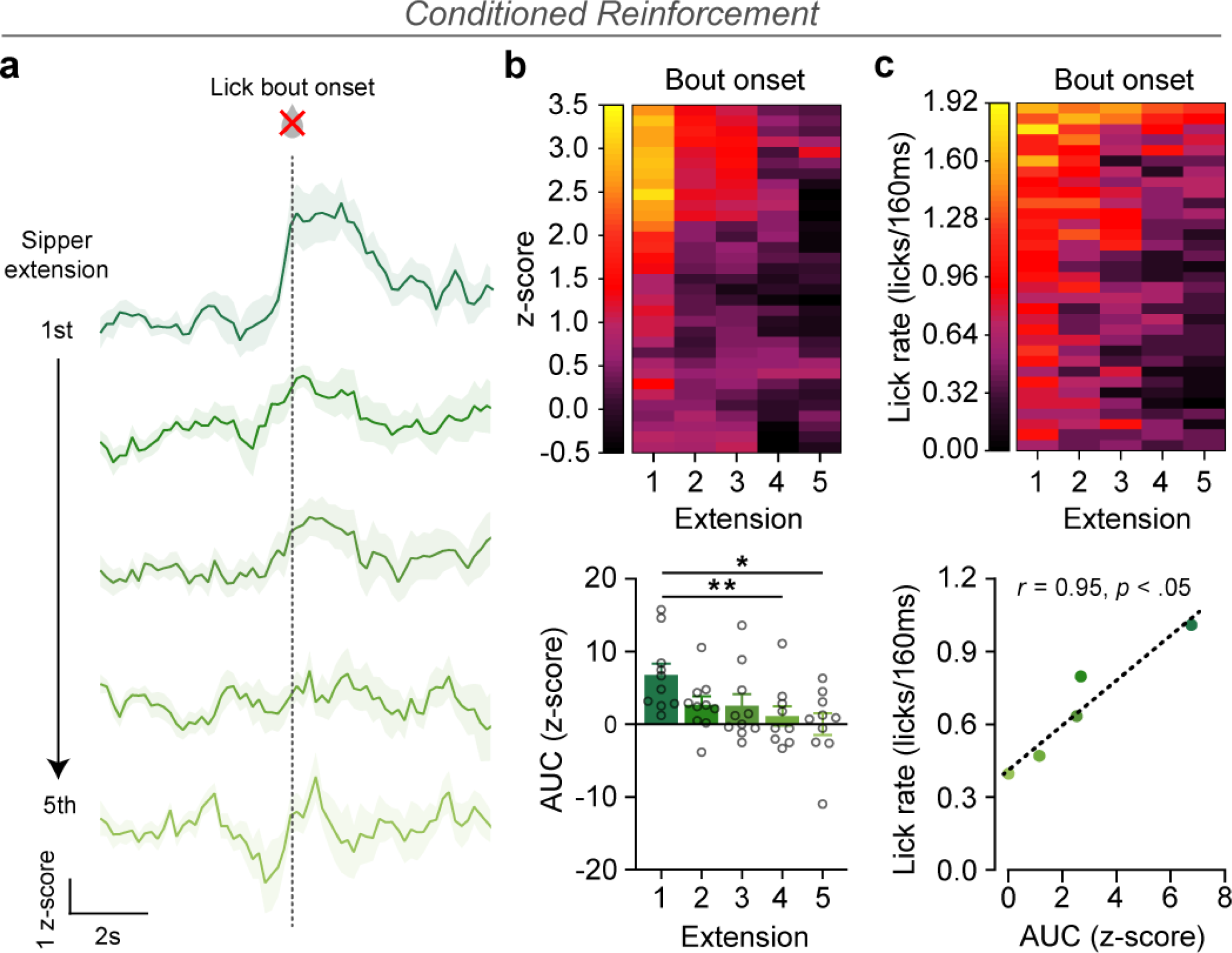
mPFC dopamine transients associated with behavioral engagement do not require an external stimulus. **(a)** Dopamine responses aligned around lick bout onset during the first 5 sipper extensions of a conditioned reinforcement session wherein the sipper tube (i.e., the conditioned reinforcer) was dry (n = 50, sampled from 10 subjects). **(b)** *Top*: Heatmap displaying dopamine activity (z-axis) averaged across animals to the lick bout onset (y-axis) for each sipper extension (x-axis). *Bottom*: Dopamine response to lick bout onset of the dry sipper tube decreased across sipper extensions (one-way repeated measures ANOVA, *F*_(2.945, 26.58)_ = 5.781, *p* = 0.0037; Tukey’s test, 1^st^ vs. 4^th^, *p* = 0.0093; 1^st^ vs. 5^th^, *p* = 0.0222; *p* > 0.05 for all other comparisons). **(c)** *Top*: Heatmap indicating the average lick rate (z-axis; 160ms bins) from lick bout onset (y-axis) for each sipper extension (x-axis). *Bottom*: There was a strong correspondence between the average lick rate and the average dopamine response aligned to licking of the dry/empty spout (Pearson’s correlation, *r* = 0.9539, *p* = 0.0118). Data represented as mean ± S.E.M. * *p* < 0.05; ** *p* < 0.01.

### mPFC Dopamine Release Signals Allocation of Attentional Resources

We next aimed to derive an explanatory construct *a posteriori* from our results thus far which would allow generation of novel, falsifiable hypotheses going forward. The results above demonstrate that mPFC dopamine release is 1) often engaged by salient stimuli but is not causally related to stimulus encoding, 2) tracks moment-to-moment changes in ongoing behavioral engagement and anticipation of proximal events, and 3) is ubiquitously modulated by novelty across scenarios, though signals are not novelty-dependent and the amplitude of modulation is modest. We reasoned that the observed dynamics are consistent with a selective attention signal, defined as the narrowing of cognitive resources towards specific aspects among all ongoing inputs^41–43^. Indeed, selective attention is modulated by external stimuli, though not causally related to stimulus encoding, is highly sensitive to novelty, and integrates internal goals with external events to guide ongoing decision-making^44–46^. Accordingly, selective attention would be predicted to be engaged during dexterous behavioral sequences, such as licking from an angled spout^47,48^, commandeered by highly salient and potentially dangerous stimuli such as tail pinch or footshock^49,50^, would not be causally altered by learning *per se*^42,51,52^, and would be expected to come online slightly prior to changes in behavioral engagement and during anticipation of arrival of an expected stimulus^53,54^. In short, selective attention is a sufficient *post hoc* explanatory construct to account for our results thus far.

In order to explicitly test this hypothesis, we experimentally manipulated attentional load in a task structure amenable to time-resolved recordings. In the discriminated operant task described above, mice were required to learn a relatively complex stimulus-response-outcome contingency. However, there was minimal attentional requirement given that the S^D^ presentation was prolonged (up to 30 seconds) and, once the contingency was learned, required only withholding responses during the S^Δ^ to achieve high performance. To specifically manipulate attentional demand without confounds related to introduction of novel stimuli, we built on the previously learned S^D^ contingency in animals that had met acquisition criteria in the discriminated operant task. Training and testing for the attentional demand task occurred in the same operant box as discriminated operant learning, featuring the same operanda, with the only physical alteration being the addition of a third lever located on the far wall from the sucrose sipper and the other two levers (referred to as the response levers). A cue light again served as the S^D^ indicating that a response on the lever below the illuminated light would be reinforced by sucrose delivery while a response on the lever below an unlighted cue would trigger a timeout. However, instead of presenting the S^D^ under a variable time schedule with a static active and inactive side, trials were self-initiated by a response on the third lever (referred to as the trial initiation lever), and the S^D^ presentation, indicating which of the two response levers would be reinforced, was pseudorandomly determined each trial (50% probability per side). Similar to the prior task, extension of a sipper containing sucrose served as the reinforcer which was delivered when a correct response was made, while a response on the alternative lever not affiliated with the S^D^ on that trial triggered a 30 second timeout period during which all three levers were retracted. During training sessions, the S^D^ was presented concomitant with a response on the trial initiation lever and remained illuminated for up to 30 seconds or until a response was made on one of the two response levers; under these conditions, mice readily learned to vary their responses from trial to trial according to the side marked by the S^D^, reaching near perfect task performance (**Extended Fig. 9**).

Finally, in the test phase of the task, a variable delay (2 – 4 second duration) was introduced between trial initiation and presentation of the S^D^, deemed the stimulus search period (**Fig. 6a**). Critically, the S^D^ was now only presented briefly (1 second duration) and the response levers remained retracted during the stimulus search and stimulus presentation periods, and extended concomitant with the termination of the S^D^. Thus, the task structure required the subject to attend during the stimulus search period but did not allow for an impulsive response prior to S^D^ presentation, by virtue of the response levers being retracted, nor did it require a working memory component given that the response could be made immediately following S^D^ offset. Congruent with demanding attention to effectively perform the task, there was a marked drop in performance under these conditions that remained well above chance (**Extended Fig. 9**). This allowed dopamine activity to be aligned to a known change point in attentional load – immediately prior to the onset of the S^D^ presentation the subject is required to allocate selective attention towards determination of the location of the imminent S^D^; during the 1 second duration of the S^D^ the stimulus location must be identified and a decision made; following the offset of the S^D^, attentional processes shift towards execution of the decision. Conforming entirely to a selective attention signal, we observed a marked reduction in mPFC dopamine activity during the stimulus search period followed by a sharp dopamine transient time-locked to the presentation of the S^D^ which resolved fully back to baseline by the end of the 1-second presentation period (**Fig. 6b,c**).

**Figure 6.**
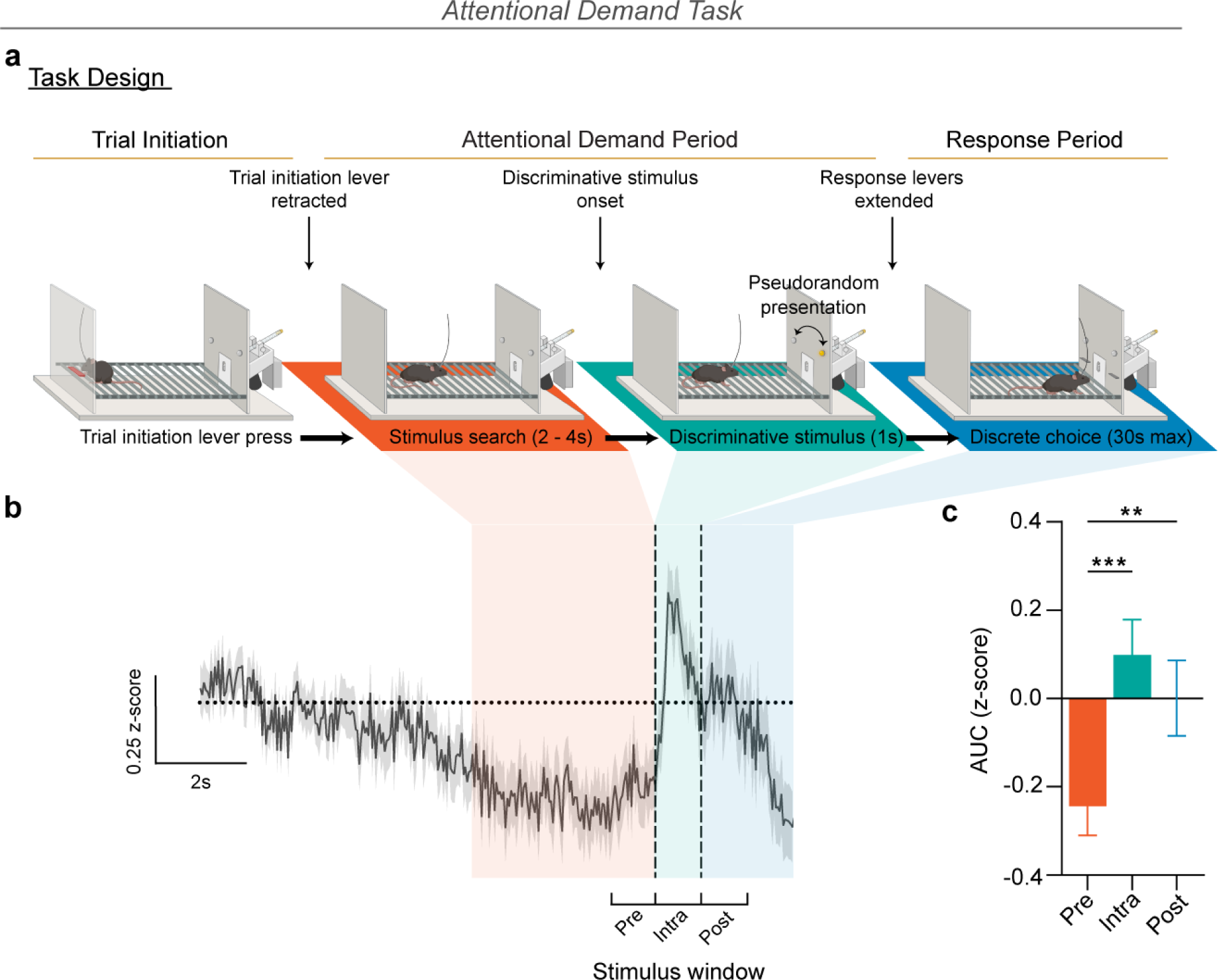
Inhibition and activation of mPFC dopamine signals attentional allocation and stimulus arrival. **(a)** Schematic of attentional demand task. A response on the trial initiation lever resulted in immediate retraction of the lever, followed by presentation of the S^D^ after a variable delay (pseudorandom, 2-4s in duration). During the delay, deemed the stimulus search period, both response levers remained retracted. The end of the stimulus search period was marked by a presentation of the S^D^, a brief (1s duration) illumination of a cue light above the correct (i.e. reinforced) lever on that trial. Concurrent with the cessation of the S^D^, both response levers were extended. A response on the lever affiliated with the S^D^ (i.e., a correct response) resulted in the extension of a sipper tube containing 10% sucrose (w/v) which was accessible for 1s commencing with first lick onset. A response on the lever unaffiliated with the S^D^ (i.e., an incorrect response) resulted in a 30s timeout period denoted by concurrent presentation of an auditory tone. After retraction of the sipper (following sucrose collection on correct response trials) or after the timeout period had elapsed (following incorrect response trials), the trial initiation lever was re-extended until another trial was initiated. **(b)** Dopamine activity aligned around S^D^ onset (n = 274 trials, sampled from 7 subjects). The vertical lines indicate S^D^ onset and offset, respectively. **(c)** Comparison of dopamine activity during three 1s epochs: 1s immediately preceding stimulus presentation (Pre), 1s concomitant with the stimulus presentation (Intra), and 1s following the offset of the stimulus (Post). Relative to the Pre-stimulus window, activity was greater during the Intra- and Post-stimulus windows (one-way repeated measures ANOVA, *F*_(1.600, 436.7)_ = 14.64, *p* < 0.0001; Tukey’s test, Pre vs. Intra, *p* < 0.0001; Pre vs. Post, *p* = 0.0065; Intra vs. Post, *p* = 0.1558). Data represented as mean ± S.E.M. ** *p* < 0.01; *** *p* < 0.001.

### Conclusions

Together, our results demonstrate that mPFC dopamine dynamics conform to a selective attention signal. In addition to the quantitative assessments detailed throughout, several observations also qualitatively lend credence to the conceptualization of mPFC dopamine subserving selective attention. First, mPFC dopamine appears to behave as a finite resource in any given period; this can be seen most clearly in the variable delay reinforcement task where the aggregate dopamine activity peristimulus is equal across trial types. Further, spontaneous transients were infrequent in a novel environment and near absent in a habituated one – the absence of ongoing activity is perhaps the most striking qualitative feature of the mPFC dopamine system, as it is in stark contrast to striatal dopamine recordings and to most circuit level processes throughout the brain where spontaneous activity is ubiquitous. The infrequency of transients outside of task-related activity and the apparent lack of redundancy of this feature in other circuits remains consistent with expectation for a selective attention signal.

Importantly, this model reconciles decades of theories but is highly congruent with prior data. For example, aversive and painful stimuli, which have been the central focus of prior theories of cortical dopamine function, are known to reflexively commandeer selective attention^49,50,55^. Similarly, competing theories have focused on higher-order executive processes but employed tasks that required selective attention^56,57^. Finally, there is a wealth of data implicating dysregulated mesocortical dopamine system function in neuropsychiatric disorders. For substance use disorder and schizophrenia, in particular, dysregulated mesocortical dopamine is posited as the causative agent driving cardinal symptomologies^58–61^. In parallel, selective attention has been directly linked to the positive symptoms of schizophrenia^62–64^ as well as narrowing of perceptual and cognitive resources towards alcohol-related stimuli in alcohol use disorder^65,66^. Thus, the proposed role of mPFC dopamine in selective attention provide a framework bridging the mesocortical dopamine system’s role in adaptive behaviors and disease.

## Author Contributions

P.R.M. and C.A.S. jointly conceived of the project and designed the experiments. P.R.M., S.O.N., C.F.F., and Z.Z.F. collected data. P.R.M. and C.A.S. developed MATLAB analysis code. P.R.M., E.K., S.O.N., and C.A.S. performed analysis. P.R.M. and C.A.S. created the figures and wrote the paper. All authors contributed to the editing of the manuscript.

## Acknowledgements

This work was supported by NIH grants R00 DA04510 (NIDA), R01 AA030115 (NIAAA), U01 AA029971 (NIAAA), Alkermes Pathways Research Award, the Brain Research Foundation, and the Whitehall Foundation (C.A.S). P.R.M. is supported by a NIH fellowship (F31 AA029626). Z.Z.F. is supported by a NIH fellowship (F31 DA056196).

## Disclosure

The authors have no conflicts to report.

## Methods

### Animals

Male C57BL/6J mice from Jackson Laboratory (Bar Harbor, ME; SN: 000664) were used for all experiments. Animals were group-housed (4-5 per cage) in a temperature- and humidity-controlled animal facility on a 12-hour reverse light-dark cycle (8 AM lights off, 8 PM lights on) with *ad libitum* access to water. Animals arrived at the facility at 8 weeks of age and were allowed to acclimate to the facility for at least 1 week before any procedures were performed and were given *ad libitum* access to chow during this period. Following acclimation and throughout all experimental procedures, chow (Picolab 5L0D, LabDiet) was given daily at slightly above caloric requirements such that a healthy adult weight was established and maintained (2.9-3 g/animal/day, corresponding to roughly 8.7 kcal^ME^/day). All experiments involving the use of animals were in accordance with NIH guidelines and approved by the Vanderbilt Institutional Animal Care and Use Committee.

### Stereotaxic Surgeries

All surgeries were conducted on mice at least 8 weeks of age using a digital small animal stereotaxic instrument (David Kopf Instruments, Tujunga, CA) under aseptic conditions and body temperature was maintained with a heating pad. Animals were anesthetized throughout surgical procedures using isoflurane (5% for induction, 1-2% for maintenance) and ophthalmic ointment was applied to both eyes to prevent corneal desiccation. A midline incision was made down the scalp and a craniotomy was performed above the injection site using a hand drill mounted to a stereotaxic arm. Unilateral (right hemisphere) 250nl injections of dLight1.2 [AAV5-hSyn-dLight1.2] (Addgene) were stereotaxically targeted to the mPFC (AP: +1.8, ML: +1.0, DV: -2.4, mm from bregma, with stereotax arm at a 10° angle away from the midline) using a beveled 33-guage microinjection needle attached to a 10µL Hamilton syringe (Neuros 1701RN, Hamilton Company). Virus was delivered at a rate of 0.1 µL per minute using a microsyringe pump (UMP3, WPI) and controller (micro2T, WPI). After 250nL were dispensed, the injection needle was left in place for at least 10 minutes. The needle was then retracted -0.05 mm towards the brain surface and allowed to rest for an additional 5 minutes before being slowly and fully retracted from the craniotomy. A chronic indwelling fiber optic probe consisting of a borosilicate optic fiber (200-µm core, 0.66 NA; Doric) housed in a metal ferrule (2.5mm diameter) was lowered to 0.1 mm above the injection site and secured to the skull with a thin layer of adhesive cement (C&B Metabond; Parkell), followed by cranioplastic cement (Ortho-Jet; Lang) mixed with black carbon powder. At the end of surgery, animals received a warmed subcutaneous injection of ketoprofen (5 mg/kg) and Ringer’s solution (∼ 1 mL), and their body temperature was maintained using a heating pad until fully recovered from anesthesia. No experiments were performed until a minimum of 6 weeks following surgery to allow for sufficient viral transduction and dLight expression.

### Ex Vivo Brain Slice Imaging

In a separate cohort of animals stereotaxically injected with dLight1.2 as previously described, following rapid decapitation, a vibrating tissue slicer (Ted Pella MicroSlicer DTK-1000N) was used to prepare 300 µM thick coronal brain sections containing the mPFC which were incubated for at least 30 minutes in aCSF oxygenated at room temperature (in mm: 126 NaCl, 2.5 KCl, 1.2 NaH_2_PO_4_, 2.4 CaCl_2_, 1.2 MgCl_2_, 25 NaHCO_3_, 11 glucose, 0.4 L-ascorbic acid, pH = 7.4). Following the incubation period, the slice was transferred to the testing chamber, which contained oxygenated aCSF at 32°C flowing at approximately 1 mL per minute. To assess the fluorescence of expressed dLight1.2 in the region in response to varying concentrations of catecholamines, slices were imaged on a custom widefield microscope setup (Cerna System, Thorlabs) with a 4x air objective (Olympus Plan Achromat Objective, 0.10 NA, 18.5 mm WD). Images were acquired at 1920 x 1080 pixel density with a pixel size of 1.26 uM (equating to 2419.2 uM by 1360.8 uM FOV size) onto a sCMOS camera (Thorlabs), thereby simultaneously capturing both the left and right hemispheres at 10 frames per second (fps). Fluorescence was averaged from a 15 second recording at each corresponding dose. Increasing concentrations (0.01, 0.1, 1, 10, and 100 µM) of dopamine (Sigma-Aldrich) or norepinephrine (Sigma-Aldrich) were systematically washed on and image stacks were taken at each dosage using both a 490 and 405 LED sequentially. At the conclusion of the experiment, image stacks were concatenated and brightness over time traces from regions of interest containing dLight1.2 expression were extracted. Traces from across the timepoints were converted to be expressed as a function of the baseline fluorescence prior to drug washes (F_0_), such that values were expressed as ΔF/F. The average ΔF/F for each dosage was calculated to determine the response to each catecholamine and in response to each LED wavelength.

### Fiber Photometry Imaging

Fiber photometry imaging was performed using a custom widefield microscope and fiber launch, similar to Kim and colleagues^67^. Briefly, 2 fiber-coupled LEDs excitation (415 and 470nm) were connected to the excitation arm of the microscope via patch cables. Each excitation source was collimated and passed through excitation filters (410nm center wavelength, 10nm FWHM and 469nm center wavelength, 35nm FWHM, respectively). The 2 beams were combined via a dichroic mirror (425nm long-pass, Thorlabs), reflected by a second dichroic (498nm long-pass, Thorlabs), and aligned to fill the back aperture of a 20x air objective (0.75 NA, Nikon MRD00205). A low autofluorescence patch cable (400µM diameter core, 0.6 NA) was held in a 3-axis translating fiber launch and adjusted such that the face of the fiber was at the focal distance of the objective (2mm). The opposite end was mated to the indwelling fiber optic cannula on the animals’ head just prior to behavioral sessions. For experiments conducted in the operant conditioning chambers, the patch cable was attached to an articulating counterbalance arm to offset the fiber weight/torque and facilitate unimpeded behavior. Time-division multiplexing of pulses of the 2 LEDs (25 Hz each, square wave) provided fluorescence excitation through the patch cable (80 ± 5 μW per channel, measured at the end of the patch cable prior to each session) output, and resulting emission was separated from the excitation light via a dichroic splitter, passed through an emission filer (525nm center wavelength, 39nm FWHM, Thorlabs) before being focused by a tube lens onto the face of a sCMOS camera imaging at 50fps (ORCA Flash, Hamamatsu). LEDs, cameras, and timestamps from behavioral equipment were synchronized by a data acquisition board (National Instruments). To pre-photobleach and minimize potential interference due to autofluorescence from the fiber optic interface, recordings we started and allowed to run for at least 60 seconds prior to beginning any behavioral task. The autofluorescence rapidly dissipates over this time, and the 60 second period is clipped out of the recording prior to performing any processing or normalization.

### Photometry Analysis

Fiber photometry data were analyzed using custom MATLAB code. A region of interest (ROI) was drawn was around the edge of the imaged fiber face and pixel intensities within the ROI were averaged per frame, resulting in two fluorescence intensity × time traces, resulting from the interleaved 405 and 470 nm excitation. The two channels were initially processed in parallel. First, each to remove photobleaching related changes fluorescence intensity, each trace was fit with a double exponential decay model, and the best fit values were then subtracted from the raw fluorescence × time trace. The residuals were then divided by the best fit values; in other words, the signals were converted to ΔF/F by the equation (*F* − *F*_0_)/ *F*_0_ where *F* is raw fluorescence at a given point in time and *F_0_* is the corresponding best-fit values from the double exponential fit. After each was ΔF/F normalized, the 410 nm trace was subtracted from the 470nm trace. Activity was then aligned to behavioral events of interest in accordance with TTL timestamps sent from the behavioral equipment which associated each event with a particular frame during the recording. Each extracted trace was then downsampled (4 times block-wise average) except for the cue-aligned traces during the attentional demand task which did not undergo downsampling. Traces were then normalized to a pre-event baseline window using a z-score transformation. For traces aligned around discrete stimuli (e.g., tailpinch, free-access sucrose), a 3 second baseline window of 5 to 2 seconds prior to stimulus onset was used. An identical baseline window was applied to traces aligned around events during continuous reinforcement, discriminated operant, and conditioned reinforcement tasks. For variable delay reinforcement and attentional demand tasks, a 3 second baseline window of 10 to 7 seconds prior to sipper extension and cue onset, respectively, was used. Z-scored traces were then averaged after trial matching when applicable to create a single trace. To quantify the magnitude of dopamine responses, peak amplitude (max value) and/or area under the curve (trapezoidal numerical integration) was calculated for each individual trace following z-score normalization and prior to averaging using a 5-second time window commencing with event onset except for traces aligned around cue onset and lever presses which were only 1 second in duration. Peaks above zero were given a positive sign while peaks below zero were given a negative sign. For area under the curve, the areas of all peaks were summed to create a net area value for each trace.

### Event Detection

For analysis of spontaneous transients in the novelty/habituation experiments, raw traces were downsampled (4 times block-wise average) and traces were smoothed using a median filter before being imported to Inscopix Data Processing Software (v1.9.2) for event detection analysis. The event detection algorithm, which selects for a fast monotonic rise in amplitude followed by an exponential decay, was then applied to the traces from each of the 20-minute sessions recorded per subject. An event threshold factor was selected and verified manually. Fluctuations in amplitude from baseline were calculated using the median absolute deviation of activity during the whole session which is a measure of statistical dispersion that is minimally affected by outliers. The average event amplitude and event rate in each session was calculated per subject.

### Lick Microstructure

Microstructural analysis of lick behavior was conducted using custom MATLAB code as previously described^68,69^. A lick bout was defined as 3 licks within 1 second of each other with the first of the 3 licks representing the bout onset. A bout was concluded when 3 seconds transpired without a lick with the final lick representing bout offset. Bout size was defined as the number of licks within a single bout while bout duration was defined as the amount of time in seconds between the onset and offset of a bout.

### Noxious Tail Pinch

Mice were placed on a cage top and allowed to acclimate for 5 minutes before commencing the tail-pinch procedure. A total of 5 tail pinches were administered by firmly pinching the base of the tail with the thumb and index finger for a duration of 3 seconds with a 60 second interval between each tail pinch onset. Mice subsequently received an intraperitoneal injection of 1 mg/kg of the D1 receptor antagonist SCH 23390 (Tocris) and were left undisturbed for 30 minutes at which time the tail-pinch procedure was repeated. SCH 23390 has been shown to act as a dLight antagonist, blocking dopamine-induced increases fluorescence^17^, and the dose was selected based on prior work demonstrating that 1 mg/kg is sufficient to saturate D1 receptors in the mPFC^70^. Animals remained tethered to the photometry patch cable throughout the entire procedure to avoid introducing variability related to changes in coupling efficiency between the indwelling and patch fibers.

### Operant reinforcement tasks

#### Overview and Apparatus (Skinner Box)

The following experiments were all performed in the dark during animals’ dark cycle inside modular 8.5” x 7.1” x 5.0” operant conditioning chambers equipped with stainless steel grid floors (Med Associates, St. Albans, Vermont). Constant 65-70 dB white noise was turned on at the start of all sessions to provide consistent ambient noise. Chambers were enclosed in sound attenuating cubicles equipped with overhead infrared cameras (Admiral 16-channel NVR; SCW) used to monitor and record each experimental session. Experiments involving exposure to discrete stimuli (e.g., novelty, free-access sucrose) were run prior to operant behavioral experiments (e.g., continuous reinforcement, discriminated operant task) except for unsignaled footshock which was run at the end of the operant experimental timeline. Accordingly, the operant chambers were progressively outfitted with modular inserts (i.e., stimuli, operanda; Med Associates) for each experiment (described below) to facilitate animals’ gradual familiarization with the operant context.

#### Novel Context

Experimentally naïve mice were placed in an unfamiliar operant chamber devoid of operanda for two 20-minute sessions conducted across consecutive days. Ambient white noise was played throughout the session as described above, but otherwise no experimental parameters were imposed.

#### Auditory Tuning Curve

Following habituation to the operant chamber, mice underwent 2 sessions across consecutive days wherein they were exposed to 6 distinct auditory tones (2.9 – 20.0 kHz in ∼2/3 octave steps, 5-second duration, 80db) presented on a variable-time (average 15 seconds) schedule. Tones were presented in a pseudorandom block design such that for every block of 6 tone presentations, each frequency was presented a single time in randomized order before a repeat stimulus was presented in the subsequent block (10 blocks in total). Tones were generated via a programmable audio generator combined with super tweeter mounted to the ceiling of the operant chamber. The dB of each frequency was tested daily, and audio card calibrated accordingly, using a sound meter attached to a microphone that was placed at the center of the operant chamber (ANL-930; Med Associates).

#### Free-access Sucrose Exposure

Mice underwent a single 20-minute magazine training session in the operant conditioning chamber during which they had unrestricted access to a sipper tube containing 10% sucrose in water (w/v). Licks were detected by a resistance lickometer grounded to the metal grid floor which was manually tested prior to all behavioral sessions involving sucrose reward. *Continuous Reinforcement:* For all subsequent tasks, operant chambers were outfitted with a retractable sipper tube flanked on both sides by retractable levers, which each had a small cue light located directly above.

During continuous reinforcement, the location of the active side was denoted by a cue light which illumuinated at the start of the session and remained on for the duration of the session. Mice were trained to respond on the active side for delivery of sucrose under continuous reinforcement (i.e. each press was reinforced). An active response resulted in the extension of a sipper tube containing 10% sucrose (w/v) through an aperture in the chamber wall, allowing access for a 10-second period following first lick contact. The inactive operandum – an identical lever on the opposite side of the sipper aperture which was distinguished only by the associated cue light remaining unlighted throughout – had no programmed consequence. The location of the active side was counterbalanced across mice. Animals completed daily 30-minute sessions until they received ≥ 75% correct responses [active responses/(active responses + inactive responses)] with a minimum of 15 correct responses in a single session.

#### Discriminated Operant Task

During the discriminated operant task, a response on the active side during a 30-second presentation of a cue light which served as a discriminative stimulus (S^D^) was deemed a correct response and resulted in the termination of the S^D^ and the extension of a sipper tube containing 10% sucrose (w/v) that was accessible for a 5-second period following first lick contact. During the S^Δ^ period, an interval period between S^D^ presentations lasting 20 – 40 seconds (average 30 seconds) wherein the cue light on the active side was not illuminated, an active response, deemed a ‘timeout response’, resulted in a 30-second timeout period signaled by the presentation of an auditory tone (12 kHz, 80 dB). Responses on either operandum during the timeout period had no consequence. Once the timeout period concluded, the auditory tone was terminated and the S^Δ^ period resumed. The location of the active side was counterbalanced across mice and responses on the inactive side had no consequence throughout the duration of the task. Animals completed daily 30-minute sessions until they received ≥ 70% correct responses [reinforcers earned/(reinforcers earned + timeouts initiated)] with a minimum of 20 reinforcers earned during each session for 2 consecutive sessions.

#### Variable Delay Reinforcement

Previously, a correct response during learning tasks resulted in the immediate extension of a sipper tube containing sucrose. For the variable delay reinforcement task, the first 3 correct responses also resulted in the immediate extension of the sipper tube. However, following these initial responses, a delay to the sipper extension (0, 2, or 5 seconds) was probabilistically introduced (equally weighted) following each correct response. Mice that acquired previous learning tasks completed a total of 3 30-minute sessions across consecutive days.

#### Conditioned Reinforcement

Mice that acquired previous learning tasks were reintroduced to the operant chambers where they underwent a single 30-minute conditioned reinforcement session wherein the task parameters were unchanged but the sipper tube (i.e., the conditioned reinforcer) was completely dry.

#### Training Task

For all subsequent tasks, operant chambers were equipped the same as the operant tasks described above but with the addition of an operandum on the side of the box opposite the retractable sipper (i.e., the trial initiation lever). During the training task, both operandum flanking the retractable sipper (i.e., the response levers) remained retracted until a trial was initiated by a response on the trial initiation lever. Upon activation, the trial initiation lever was immediately retracted, and the S^D^ was pseudorandomly presented above one of the response levers which were extended concomitant with S^D^ onset. Mice were then given a 30-second period to respond throughout which the cue light representing the S^D^ remained continuously illuminated. A correct response occurred when a response was made on the lever situated below the S^D^ resulting in the extension of a sipper tube containing 10% sucrose (w/v) that remained accessible for a 1-second period commencing with first lick onset. A timeout response was made when the lever situated below the non-illuminated cue light was activated. This resulted in a 30-second timeout period signaled by the simultaneous presentation of an auditory tone, termination of the S^D^, and the retraction of both response levers. Following the completion of each trial, the trial initiation lever was re-extended to allow for the initiation of the next trial. Mice completed daily 30-minute sessions until they achieved ≥ 90% correct trials (correct trials/total trials) with a minimum of 40 correct trials in a single session.

#### Attentional Demand Task

After the training task, animals completed 4 total 30-minute sessions across consecutive days of a modified version of the training task featuring 3 changes to the previous task design to increase attentional demand. First, a variable delay period was introduced (2 – 4 seconds) between the initiation of a trial and the onset of the illuminated cue light (i.e. the S^D^). Second, the duration of the S^D^ was reduced from 30 seconds to 1 second. Third, the response levers remained retracted following trial initiation and were only extended upon the offset of the S^D^. Following termination of the S^D^, animals had a 30 second period to respond during which a response on the lever affiliated with the S^D^ resulted in the extension of the sipper tube for 1 second commencing with lick onset. A response on the lever not affiliated with the S^D^ resulted in a 30-second timeout period.

#### Unsignaled Footshock

During a single session, mice received a total of 12 footshocks (1 second duration) comprised of triplicate series of 4 footshocks of increasing intensity (0.2mA→0.4mA→0.6mA→0.8mA). Shocks were delivered non-contingently with a variable 30-second inter-stimulus interval via a computer-controlled constant current stimulator. Prior to each session, shock output was systematically tested across the grid floor of the operant chamber with an amp meter (ENV-420; Med Associates) to ensure that amperage was consistent and accurate across all grid floor bars.

### Histology

Mice were deeply anaesthetized before being transcardially perfused with 10 mL of 1x PBS solution followed by 10 mL of cold 4% PFA in 1x PBS. Animals were then rapidly decapitated, and the brain was extracted and stored at 4 °C in a vial containing 4% PFA for at least 48 hours. Prior to slicing, brains were transferred to a 30% sucrose solution in 1x PBS and kept at 4 °C until brains sank to the bottom of the vial. Upon sinking, brains were sectioned at 40µm on a freezing sliding microtome (HM 430; Thermo Fisher Scientific). Prior to each step of immunohistological processing, sections underwent 4x 10 min washes in 1x PBS. Sections were immunohistochemically stained with an anti-GFP antibody (chicken anti-GFP, 1:2000; Aves Labs) and stored overnight at room temperature. Slices were then incubated with a secondary antibody (donkey anti-chicken AlexaFluor 488, 1:500) before being covered and stored °at 4 °C overnight. To achieve fluorescent staining of nuclei, sections were incubated in DAPI (Thermo Fisher Scientific) for 5 minutes and then mounted on glass microscope slides (Thermo Fisher Scientific).

### Statistics

All statistical analyses were performed using GraphPad Prism V10 (GraphPad Software, Boston, Massachusetts). Comparisons across 2 or more conditions were made using nested one-way ANOVAs followed by Tukey’s multiple comparison test. Comparisons across 2 time points were performed using a paired samples t-test while comparisons across 3 or more time points were performed using a repeated measures one-way ANOVA followed by Tukey’s multiple comparison test. For analyses involving comparisons of multiple conditions across time points, a two-way repeated measures ANOVA was used followed by Šídák’s multiple comparison test. All tests were two-sided and p values < 0.05 were considered to be statistically significant.

## Notes

### Competing Interest Statement

The authors have declared no competing interest.

## References

1. Goldman-Rakic, P. S. Dopamine-mediated mechanisms of the prefrontal cortex. Semin. Neurosci. 4, 149– 159 (1992).

2. Weele, C. M. V., Siciliano, C. A. & Tye, K. M. Dopamine tunes prefrontal outputs to orchestrate aversive processing. Brain Res. 1713, 16–31 (2019).

3. Seamans, J. K. & Yang, C. R. The principal features and mechanisms of dopamine modulation in the prefrontal cortex. Prog. Neurobiol. 74, 1–58 (2004).

4. Robbins, T. W. Chemistry of the mind: neurochemical modulation of prefrontal cortical function. J. Comp. Neurol. 493, 140–146 (2005).

5. Fox, M. E. & Wightman, R. M. Contrasting Regulation of Catecholamine Neurotransmission in the Behaving Brain: Pharmacological Insights from an Electrochemical Perspective. Pharmacol. Rev. 69, 12– 32 (2017).

6. Thierry, A. M., Tassin, J. P., Blanc, G. & Glowinski, J. Selective activation of mesocortical DA system by stress. Nature 263, 242–244 (1976).

7. Mantz, J., Thierry, A. M. & Glowinski, J. Effect of noxious tail pinch on the discharge rate of mesocortical and mesolimbic dopamine neurons: selective activation of the mesocortical system. Brain Res. 476, 377– 381 (1989).

8. Roth, R. H., Tam, S. Y., Ida, Y., Yang, J. X. & Deutch, A. Y. Stress and the mesocorticolimbic dopamine systems. Ann. N. Y. Acad. Sci. 537, 138–147 (1988).

9. Valzelli, L. & Garattini, S. Biogenic amines in discrete brain areas after treatment with monoamineoxidase inhibitors. J. Neurochem. 15, 259–261 (1968).

10. Thierry, A. M., Blanc, G., Sobel, A., Stinus, L. & Glowinski, J. Dopaminergic terminals in the rat cortex. Science 182, 499–501 (1973).

11. Thierry, A. M., Stinus, L., Blanc, G. & Glowinski, J. Some evidence for the existence of dopaminergic neurons in the rat cortex. Brain Res. 50, 230–234 (1973).

12. Miner, L. H., Schroeter, S., Blakely, R. D. & Sesack, S. R. Ultrastructural localization of the norepinephrine transporter in superficial and deep layers of the rat prelimbic prefrontal cortex and its spatial relationship to probable dopamine terminals. J. Comp. Neurol. 466, 478–494 (2003).

13. Sesack, S. R. & Pickel, V. M. Dual ultrastructural localization of enkephalin and tyrosine hydroxylase immunoreactivity in the rat ventral tegmental area: multiple substrates for opiate-dopamine interactions. J. Neurosci. 12, 1335–1350 (1992).

14. Shnitko, T. A., Kennerly, L. C., Spear, L. P. & Robinson, D. L. Ethanol reduces evoked dopamine release and slows clearance in the rat medial prefrontal cortex. Alcohol. Clin. Exp. Res. 38, 2969–2977 (2014).

15. Garris, P. A., Collins, L. B., Jones, S. R. & Wightman, R. M. Evoked extracellular dopamine in vivo in the medial prefrontal cortex. J. Neurochem. 61, 637–647 (1993).

16. Vander Weele, C. M., et al. Dopamine enhances signal-to-noise ratio in cortical-brainstem encoding of aversive stimuli. Nature 563, 397–401 (2018).

17. Patriarchi, T. et al. Ultrafast neuronal imaging of dopamine dynamics with designed genetically encoded sensors. Science 360, (2018).

18. Patriarchi, T. et al. An expanded palette of dopamine sensors for multiplex imaging in vivo. Nat. Methods 17, 1147–1155 (2020).

19. Lammel, S. et al. Input-specific control of reward and aversion in the ventral tegmental area. Nature 491, 212–217 (2012).

20. Lammel, S., Ion, D. I., Roeper, J. & Malenka, R. C. Projection-specific modulation of dopamine neuron synapses by aversive and rewarding stimuli. Neuron 70, 855–862 (2011).

21. Deutch, A. Y. & Roth, R. H. Chapter 19 The determinants of stress-induced activation of the prefrontal cortical dopamine system. in Progress in Brain Research (eds. Uylings, H. B. M., Van Eden, C. G., De Bruin, J. P. C., Corner, M. A. & Feenstra, M. G. P.) vol. 85 367–403 (Elsevier, 1991).

22. Reinhard, J. F., Jr, Bannon, M. J. & Roth, R. H. Acceleration by stress of dopamine synthesis and metabolism in prefrontal cortex: antagonism by diazepam. Naunyn. Schmiedebergs. Arch. Pharmacol. 318, 374–377 (1982).

23. Deutch, A. Y., Tam, S. Y. & Roth, R. H. Footshock and conditioned stress increase 3,4-dihydroxyphenylacetic acid (DOPAC) in the ventral tegmental area but not substantia nigra. Brain Res. 333, 143–146 (1985).

24. Fadda, F. et al. Stress-induced increase in 3,4-dihydroxyphenylacetic acid (DOPAC) levels in the cerebral cortex and in n. accumbens: Reversal by diazepam. Life Sci. 23, 2219–2224 (1978).

25. Lavielle, S. et al. Blockade by benzodiazepines of the selective high increase in dopamine turnover induced by stress in mesocortical dopaminergic neurons of the rat. Brain Res. 168, 585–594 (1979).

26. Bassareo, V., Tanda, G., Petromilli, P., Giua, C. & Di Chiara, G. Non-psychostimulant drugs of abuse and anxiogenic drugs activate with differential selectivity dopamine transmission in the nucleus accumbens and in the medial prefrontal cortex of the rat. Psychopharmacology 124, 293–299 (1996).

27. Bassareo, V., De Luca, M. A. & Di Chiara, G. Differential Expression of Motivational Stimulus Properties by Dopamine in Nucleus Accumbens Shell versus Core and Prefrontal Cortex. J. Neurosci. 22, 4709–4719 (2002).

28. Hernandez, L. & Hoebel, B. G. Feeding can enhance dopamine turnover in the prefrontal cortex. Brain Res. Bull. 25, 975–979 (1990).

29. St Onge, J. R., Ahn, S., Phillips, A. G. & Floresco, S. B. Dynamic fluctuations in dopamine efflux in the prefrontal cortex and nucleus accumbens during risk-based decision making. J. Neurosci. 32, 16880– 16891 (2012).

30. Floresco, S. B. & Magyar, O. Mesocortical dopamine modulation of executive functions: beyond working memory. Psychopharmacology 188, 567–585 (2006).

31. Seamans, J. K. & Robbins, T. W. Dopamine Modulation of the Prefrontal Cortex and Cognitive Function. In The Dopamine Receptors (ed. Neve, K. A.) 373–398 (Humana Press, 2010).

32. Lapish, C. C., Kroener, S., Durstewitz, D., Lavin, A. & Seamans, J. K. The ability of the mesocortical dopamine system to operate in distinct temporal modes. Psychopharmacology 191, 609–625 (2007).

33. Toth, B. A., Chang, K. S., Fechtali, S. & Burgess, C. R. Dopamine release in the nucleus accumbens promotes REM sleep and cataplexy. iScience 26, 107613 (2023).

34. *Kappa Opioid Receptors Negatively Regulate Real Time Spontaneous Dopamine Signals by Reducing Release and Increasing Uptake*.

35. Lammel, S. et al. Unique properties of mesoprefrontal neurons within a dual mesocorticolimbic dopamine system. Neuron 57, 760–773 (2008).

36. Kutlu, M. G. et al. Dopamine release in the nucleus accumbens core signals perceived saliency. Curr. Biol. 31, 4748–4761.e8 (2021).

37. Flagel, S. B. et al. A selective role for dopamine in stimulus-reward learning. Nature 469, 53–57 (2011).

38. Day, J. J., Roitman, M. F., Wightman, R. M. & Carelli, R. M. Associative learning mediates dynamic shifts in dopamine signaling in the nucleus accumbens. Nat. Neurosci. 10, 1020–1028 (2007).

39. Phillips, P. E. M., Stuber, G. D., Heien, M. L. A. V., Wightman, R. M. & Carelli, R. M. Subsecond dopamine release promotes cocaine seeking. Nature 422, 614–618 (2003).

40. Keiflin, R. & Janak, P. H. Dopamine prediction errors in reward learning and addiction: From theory to neural circuitry. Neuron 88, 247–263 (2015).

41. Hahn, B. et al. Divided versus selective attention: Evidence for common processing mechanisms. Brain Res. 1215, 137–146 (2008).

42. Johnston, W. A. & Dark, V. J. Selective attention. Annu. Rev. Psychol. 37, 43–75 (1986).

43. Broadbent, D. E. Perception and communication. (Pergamon Press, 1958).

44. Anderson, B. A. A value-driven mechanism of attentional selection. J. Vis. 13, (2013).

45. van Ede, F., Board, A. G. & Nobre, A. C. Goal-directed and stimulus-driven selection of internal representations. Proc. Natl. Acad. Sci. U. S. A. 117, 24590–24598 (2020).

46. Chun, M. M., Golomb, J. D. & Turk-Browne, N. B. A taxonomy of external and internal attention. Annu. Rev. Psychol. 62, 73–101 (2011).

47. Vajnerová, O., Bielavská, E., Jiruska, P. & Brozek, G. Level of vigilance influences licking frequency in rats. Physiol. Res. 52, 243–249 (2003).

48. Weijnen, J. A. Licking behavior in the rat: measurement and situational control of licking frequency. Neurosci. Biobehav. Rev. 22, 751–760 (1998).

49. Klein, Z., Ginat-Frolich, R., Barry, T. J. & Shechner, T. Effects of increased attention allocation to threat and safety stimuli on fear extinction and its recall. J. Behav. Ther. Exp. Psychiatry 72, 101640 (2021).

50. Richards, H. J., Benson, V., Donnelly, N. & Hadwin, J. A. Exploring the function of selective attention and hypervigilance for threat in anxiety. Clin. Psychol. Rev. 34, 1–13 (2014).

51. Krauzlis, R. J., Bogadhi, A. R., Herman, J. P. & Bollimunta, A. Selective attention without a neocortex. Cortex 102, 161–175 (2018).

52. Kruschke, J. K. Models of attentional learning. in Formal Approaches in Categorization (eds. Pothos, E. M. & Wills, A. J.) 120–152 (Cambridge University Press, 2011).

53. Driver, J. & Frackowiak, R. S. Neurobiological measures of human selective attention. Neuropsychologia 39, 1257–1262 (2001).

54. Watamaniuk, S. N. J. & Heinen, S. J. Allocation of attention during pursuit of large objects is no different than during fixation. J. Vis. 15, 9 (2015).

55. Kim, H. & Anderson, B. A. How does the attention system learn from aversive outcomes? Emotion 21, 898–903 (2021).

56. Hanania, R. & Smith, L. B. Selective attention and attention switching: towards a unified developmental approach. Dev. Sci. 13, 622–635 (2010).

57. Goldman-Rakic, P. S. The cortical dopamine system: role in memory and cognition. Adv. Pharmacol. 42, 707–711 (1998).

58. 58. The dopamine hypothesis of schizophrenia: focus on the dopamine receptor. Am. J. Psychiatry 133, 197–202 (1976).

59. Howes, O. D. & Kapur, S. The dopamine hypothesis of schizophrenia: version III--the final common pathway. Schizophr. Bull. 35, 549–562 (2009).

60. Abernathy, K., Chandler, L. J. & Woodward, J. J. Alcohol and the Prefrontal Cortex. in International Review of Neurobiology (eds. Reilly, M. T. & Lovinger, D. M.) vol. 91 289–320 (Academic Press, 2010).

61. Volkow, N. D. & Morales, M. The brain on drugs: From reward to addiction. Cell 162, 712–725 (2015).

62. Morris, R., Griffiths, O., Le Pelley, M. E. & Weickert, T. W. Attention to irrelevant cues is related to positive symptoms in schizophrenia. Schizophr. Bull. 39, 575–582 (2013).

63. Egeland, J. et al. Attention profile in schizophrenia compared with depression: differential effects of processing speed, selective attention and vigilance. Acta Psychiatr. Scand. 108, 276–284 (2003).

64. Oltmanns, T. F. Selective attention in schizophrenic and manic psychoses: The effect of distraction on information processing. J. Abnorm. Psychol. 87, 212–225 (1978).

65. Cordovil De Sousa Uva, M., et al. Distinct effects of protracted withdrawal on affect, craving, selective attention and executive functions among alcohol-dependent patients. Alcohol Alcohol 45, 241–246 (2010).

66. Harvey, A. J. When alcohol narrows the field of focal attention. Q. J. Exp. Psychol. (Hove) 69, 669–677 (2016).

67. Kim, C. K. et al. Simultaneous fast measurement of circuit dynamics at multiple sites across the mammalian brain. Nat. Methods 13, 325–8 (2016).

68. Siciliano, C. A. et al. A cortical-brainstem circuit predicts and governs compulsive alcohol drinking. Science 366, 1008–1012 (2019).

69. Brown, A. R. et al. Structured tracking of alcohol reinforcement (STAR) for basic and translational alcohol research. Mol. Psychiatry 28, 1585–1598 (2023).

70. Neisewander, J. L., Fuchs, R. A., O’Dell, L. E. & Khroyan, T. V. Effects of SCH-23390 on dopamine D1 receptor occupancy and locomotion produced by intraaccumbens cocaine infusion. Synapse 30, 194–204 (1998).

