## Supplemental Figures & Legends for "Medial prefrontal dopamine dynamics reflect allocation of selective attention"

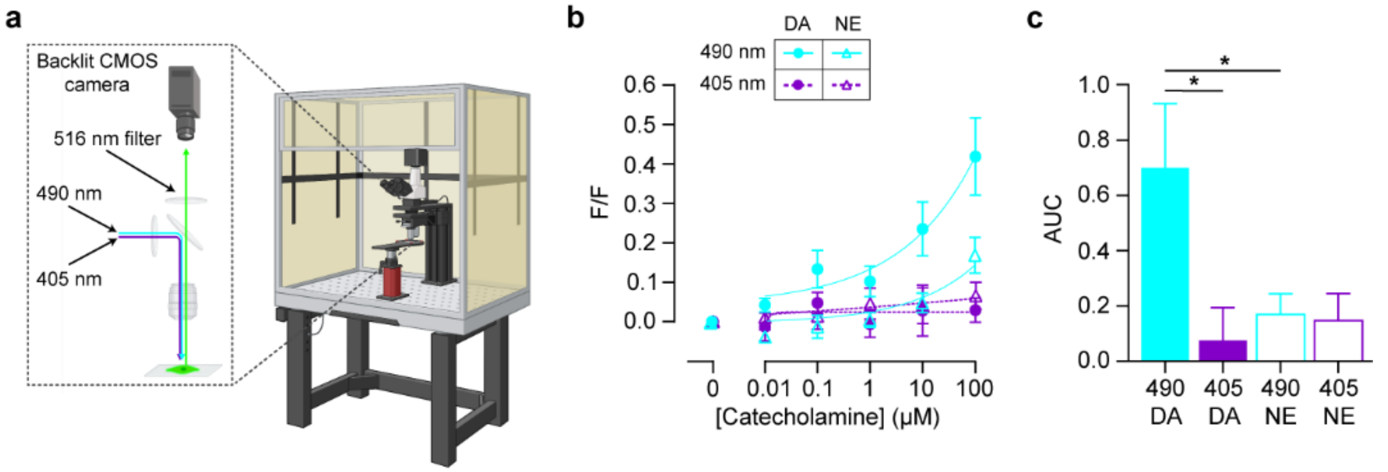


**Extended Data Figure 1. Catecholaminergic selectivity of dLight1.2.** To determine the selectivity of dLight1.2 for dopamine over norepinephrine, mouse brain slices containing the medial prefrontal cortex (mPFC) of animals previously stereotaxically injected with dLight1.2 were imaged *ex vivo*. Cumulative concentrations (0.01 to 100 µM) of dopamine (DA) or norepinephrine (NE) were added to the aCSF perfusate (separate slices), and timelapse images were acquired at 3-minute intervals using either 490 nm or 405 nm excitation. Pixel intensities across the field of view were averaged, and brightness at each timepoint (F) was expressed as F/F $[(F/F_0)]$ where $F_0$ was the mean field of view intensity prior to wash on. **(b)** Concentration response curves for DA and NE using 490 and 405 nm excitation. **(c)** Area under the curve (AUC) analysis demonstrates robust increases in fluorescence for DA using 490 nm excitation compared to NE and 405 nm excitation (one-way ANOVA, $F_{(3,24)} = 3.695$, $p = 0.0256$; planned Šídák's multiple comparisons test, 490 DA vs. 490 NE, $p = 0.0469$; 490 DA vs. 405 DA, $p = 0.0281$; 490 NE vs. 405 NE, $p = 0.9995$). Data represented as mean ± S.E.M. * $p < 0.05$.

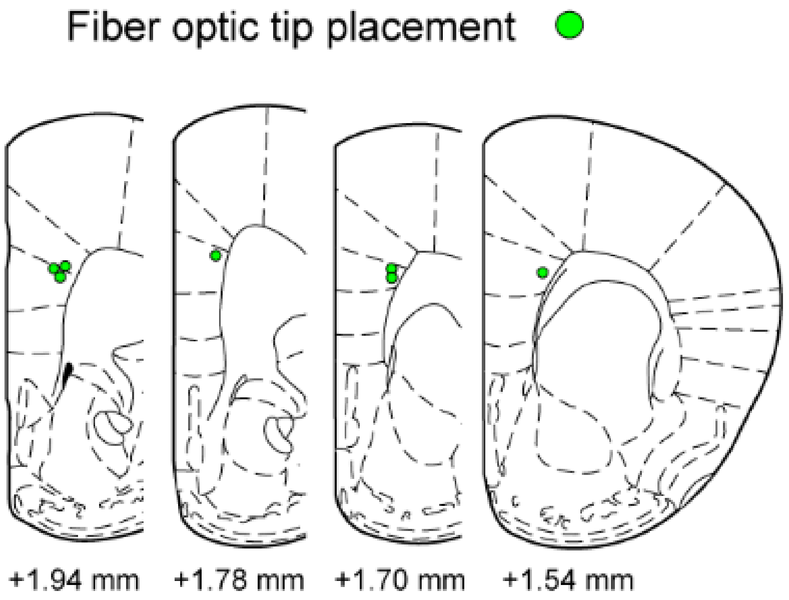


**Extended Data Figure 2. Site of dLight1.2 fiber photometry recordings.** Schematic of fiber optic placements (green) in the right hemisphere of the mPFC of experimental animals. Coordinates represent distance relative to bregma. Atlas images were reproduced from Paxinos & Franklin (2001) 2nd edition.

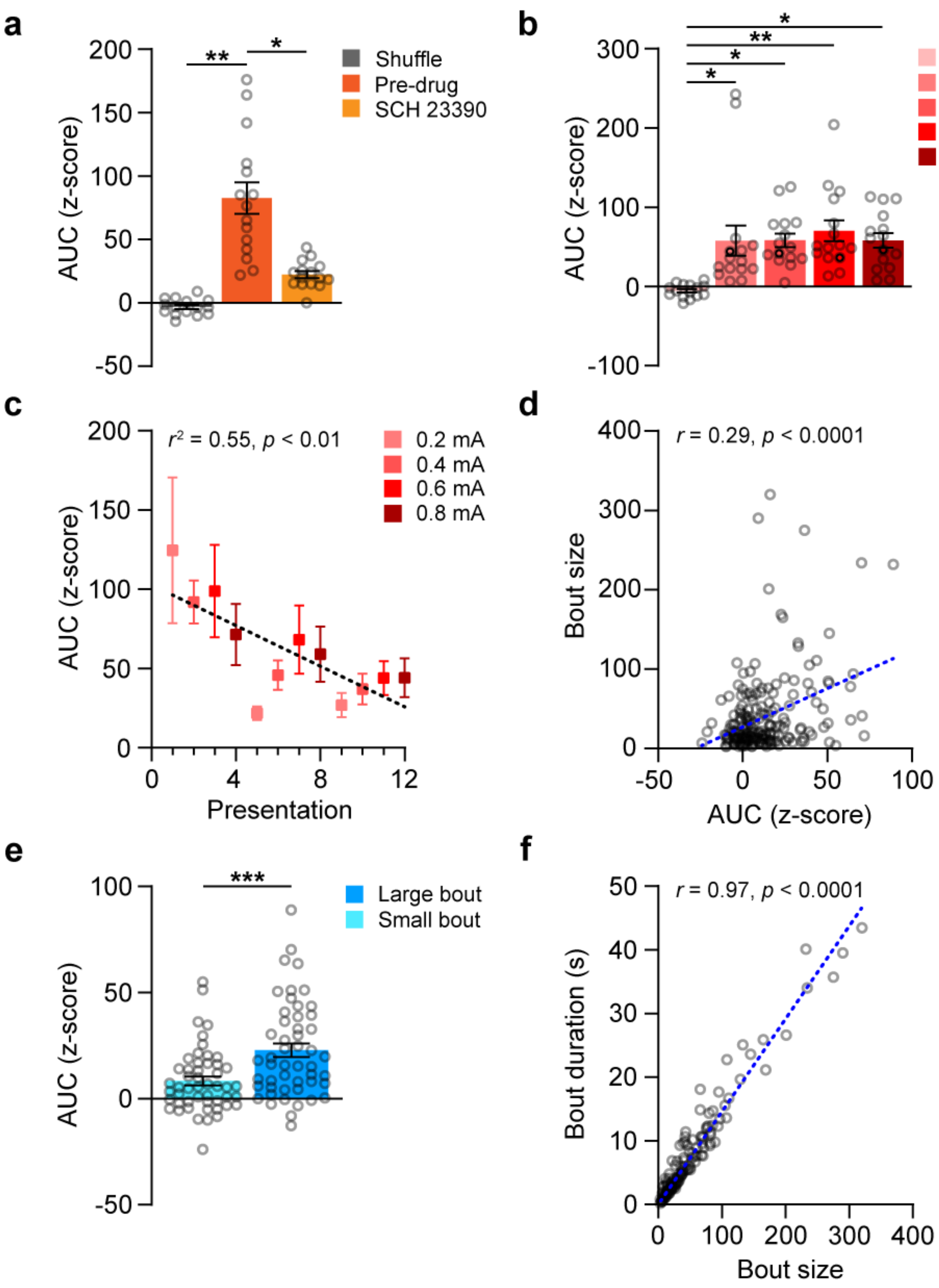
a
200
150
100
50
0
-50
AUC (z-score)
**
*
Shuffle
Pre-drug
SCH 23390
b
300
200
100
0
-100
AUC (z-score)
*
**
*
*
Shuffle
0.2 mA
0.4 mA
0.6 mA
0.8 mA
c
r² = 0.55, p < 0.01
0.2 mA
0.4 mA
0.6 mA
0.8 mA
AUC (z-score)
0
4
8
12
Presentation
d
r = 0.29, p < 0.0001
400
300
200
100
0
Bout size
-50
0
50
100
AUC (z-score)
e
***
Large bout
Small bout
AUC (z-score)
f
r = 0.97, p < 0.0001
50
40
30
20
10
0
Bout duration (s)
0
100
200
300
400
Bout size

**Extended Data Figure 3. mPFC dopamine activity is valence independent. (a)** Congruent with peak amplitude analyses presented in Figure 1, AUC of the tail pinch-evoked mPFC dopamine response was greater than the shuffled time alignment and was attenuated by systemic delivery of the dLight antagonist SCH 23390 (1 mg/kg, i.p.) (nested ANOVA, $F_{(2,6)} = 14.14$, $p = 0.0054$; Tukey's test, Shuffle vs. Pre-drug, $p = 0.0050$; Shuffle vs. SCH 23390, $p = 0.3421$; Pre-drug vs. SCH 23390, $p = 0.0251$). **(b)** The AUC of footshock-evoked responses were greater than the shuffled time alignment but did not differ as a function of amperage (nested ANOVA, $F_{(4,20)} = 5.328$, $p = 0.0044$; Tukey's test, Shuffle vs. 0.2mA, $p = 0.0207$; Shuffle vs. 0.4mA, $p = 0.0197$; Shuffle vs. 0.6mA, $p = 0.0046$; Shuffle vs. 0.8mA, $p = 0.0197$; $p > 0.05$ for all other comparisons). **(c)** The AUC of footshock-evoked dopamine responses decreased linearly as a function of presentation order regardless of amperage (simple linear regression, $r^2 = 0.5528$, $F_{(1,10)} = 12.36$, $p = 0.0056$). **(d)** The size of individual lick bouts was correlated with the AUC of the dopamine response (Spearman's correlation, $r = 0.2879$, $p < 0.0001$). **(e)** The AUC of the dopamine response associated with large (upper quartile) bout sizes was greater than the response during small (lower quartile) bouts (Mann-Whitney U test, $U = 739$, $p = 0.0004$). **(f)** There was a near-exact correspondence between the number of licks in a bout and the lick bout duration (Pearson's correlation, $r = 0.9742$, $p < 0.0001$), thus for later analysis only bout size is presented for brevity. Data represented as mean $\pm$ S.E.M. * $p < 0.05$; ** $p < 0.01$; *** $p < 0.001$.

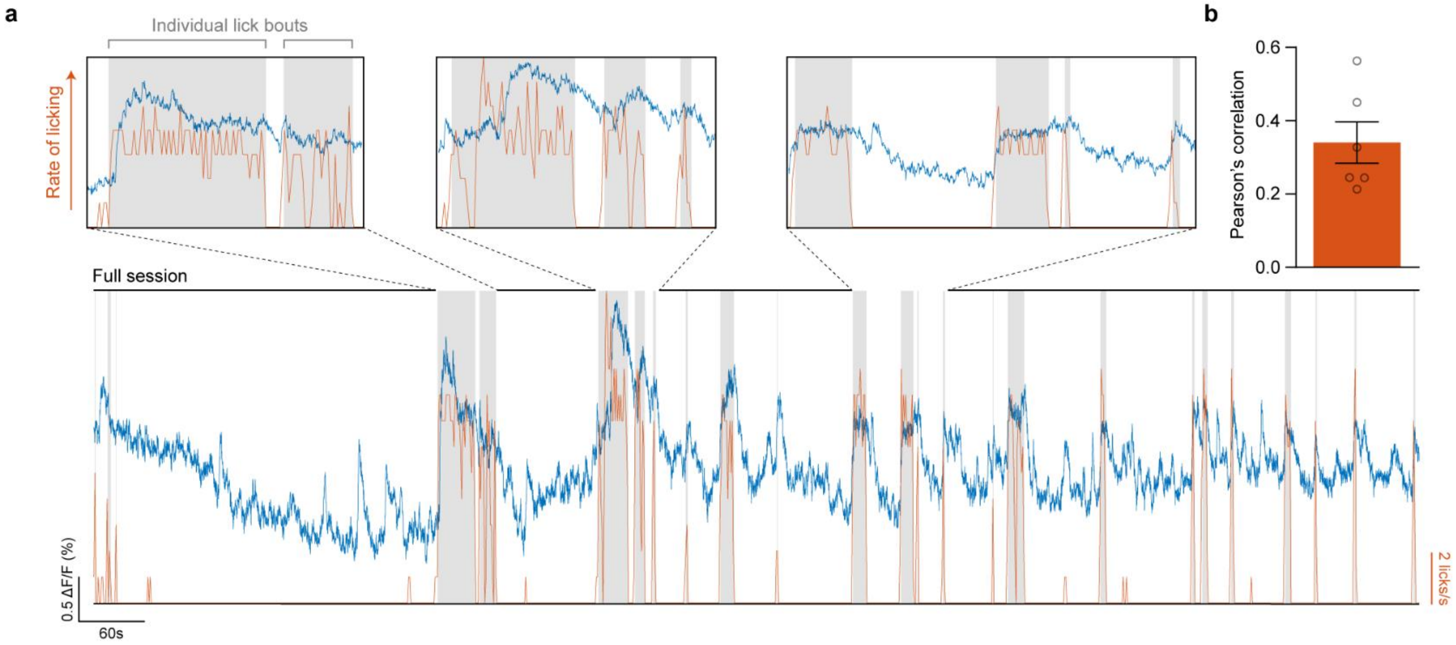


**Extended Data Figure 4. Sub-second fluctuations in mPFC dopamine scale with ongoing consummatory engagement. (a)** mPFC dopamine activity from a single animal during a 20-minute session wherein animals were given open access to a lick spout containing sucrose solution (10% w/v) illustrating lick rate (orange) and within-bout periods (gray) overlaid with dLight1.2 fluorescence (blue). Close-ups of the data are presented in the top row for clear visualization. **(b)** Pearson's correlations from 6 subjects demonstrating correspondence between dopamine activity and the rate of licking throughout sucrose exposure session. Data in panel b are represented as mean ± S.E.M.

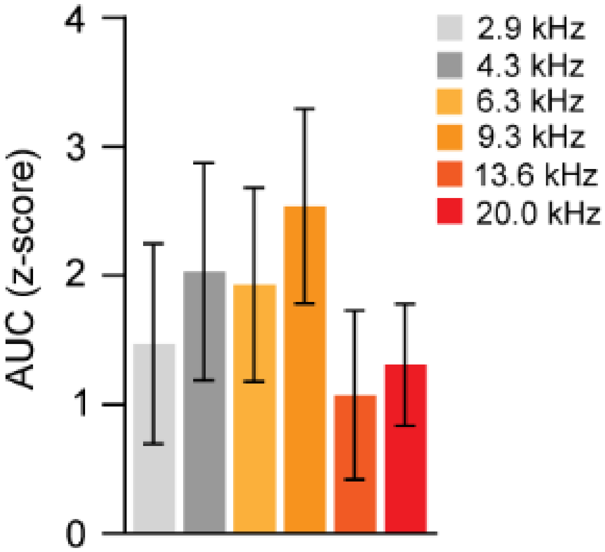


**Extended Data Figure 5. mPFC dopamine response to auditory tones is not frequency dependent.** Across both sessions of auditory tone exposure there was no difference in tone-evoked dopamine response as a function of frequency (nested ANOVA, $F_{(5,24)} = 0.4227$, $p = 0.8283$). Data represented as mean $\pm$ S.E.M.

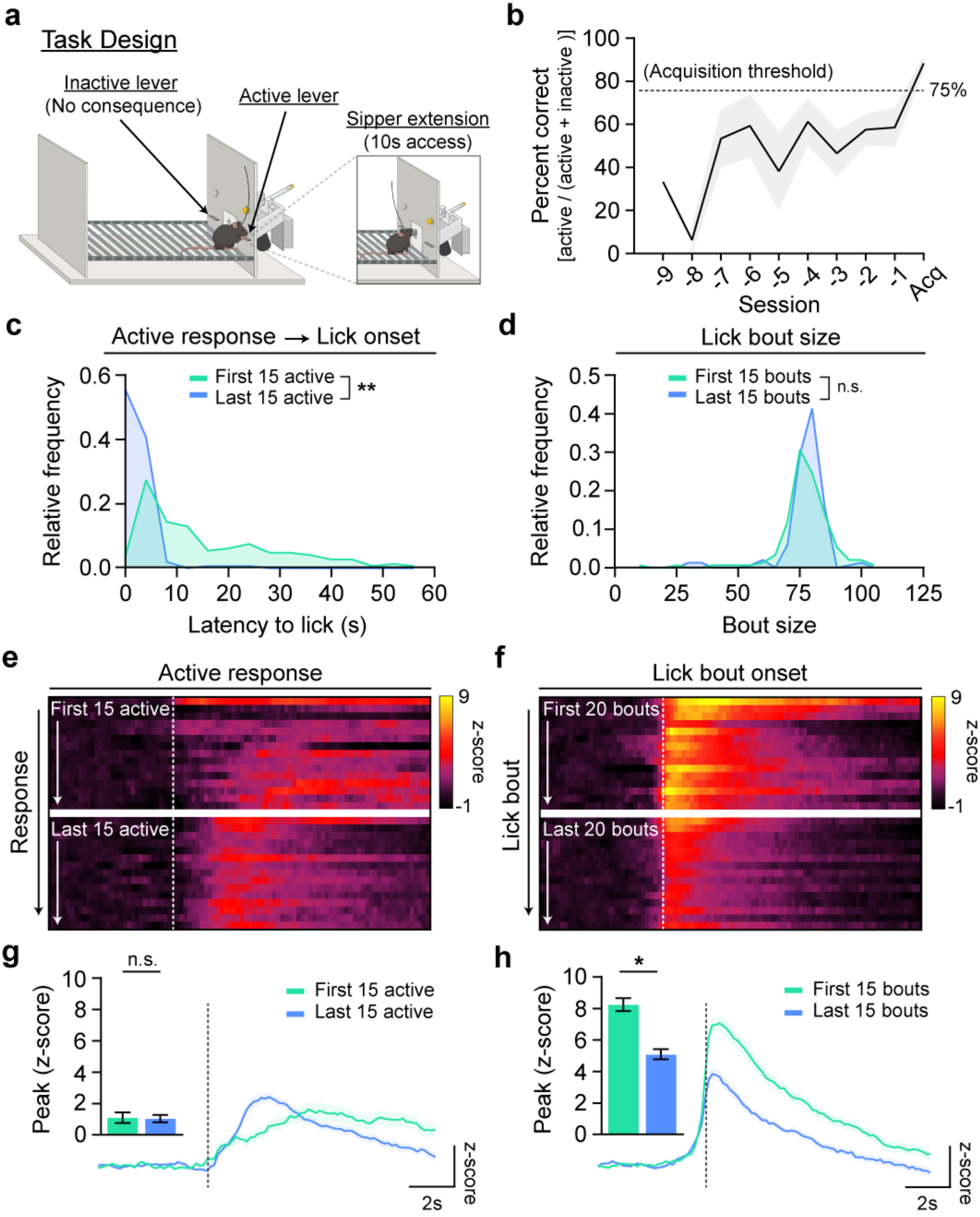
a
Task Design
Inactive lever
(No consequence)
Active lever
Sipper extension
(10s access)
b
Percent correct
[active / (active + inactive )]
(Acquisition threshold)
75%
Session
Acq
c
Active response → Lick onset
First 15 active
Last 15 active
**
Relative frequency
Latency to lick (s)
d
Lick bout size
First 15 bouts
Last 15 bouts
n.s.
Relative frequency
Bout size
e
Active response
First 15 active
Last 15 active
Response
z-score
f
Lick bout onset
First 20 bouts
Last 20 bouts
Lick bout
z-score
g
n.s.
Peak (z-score)
First 15 active
Last 15 active
2s
z-score
h
*
Peak (z-score)
First 15 bouts
Last 15 bouts
2s
z-score

**Extended Data Figure 6. Minimal changes in mPFC dopamine activity across acquisition of a positive reinforcement contingency. (a)** Schematic of continuous reinforcement task (i.e. fixed-ratio 1). A cue light was illuminated above the active operandum throughout the duration of each session. Each response on the active operandum was reinforced by the extension of a sipper tube containing 10% sucrose (w/v) for a 10 second period commencing with first lick contact, while responses on the inactive operandum had no consequence. **(b)** Animals were tested until performance reached ≥ 75% active responses [active responses / (active responses + inactive responses)] while attaining a minimum of 15 active responses in a single session. **(c,d)** Comparison of behavioral measures during the first 15 active responses during the pre-acquisition period and the first 15 active responses during the day of acquisition (Acq). **(c)** By the final session, animals displayed lower latencies to initiate a lick bout following an active response (nested ANOVA, $F_{(1,18)} = 64.50$, $p < 0.0001$) **(d)** but no difference in the number of licks per bout (nested ANOVA, $F_{(1,18)} = 0.0088$, $p = 0.9263$). **(e)** Heatmap displaying dopamine activity (z-axis) surrounding active responses (x-axis) averaged across animals for each of the first 15 active responses (y-axis) during pre-acquisition and the acquisition session. **(f)** Heatmap displaying dopamine activity (z-axis) surrounding lick bout onset (x-axis) averaged across animals for each of the first 15 lick bouts (y-axis) during pre-acquisition and the acquisition session. **(g)** Averaged dopamine traces (n = 150 events each first/last, sampled from 10 subjects). Vertical line indicates triggering of an active response. *Inset:* There was no change in dopamine activity at the time of an active response before vs after learning (nested ANOVA, $F_{(1,18)} = 0.0134$, $p = 0.9092$). **(h)** Averaged dopamine traces (n = 150 events each first/last, sampled from 10 subjects). Vertical line indicates first lick contact following sipper extension. *Inset:* The magnitude of the dopamine response following lick bout onset decreased after learning (nested ANOVA, $F_{(1,18)} = 6.631$, $p$ 0.0191). Data represented as mean ± S.E.M. * $p < 0.05$; ** $p < 0.01$.

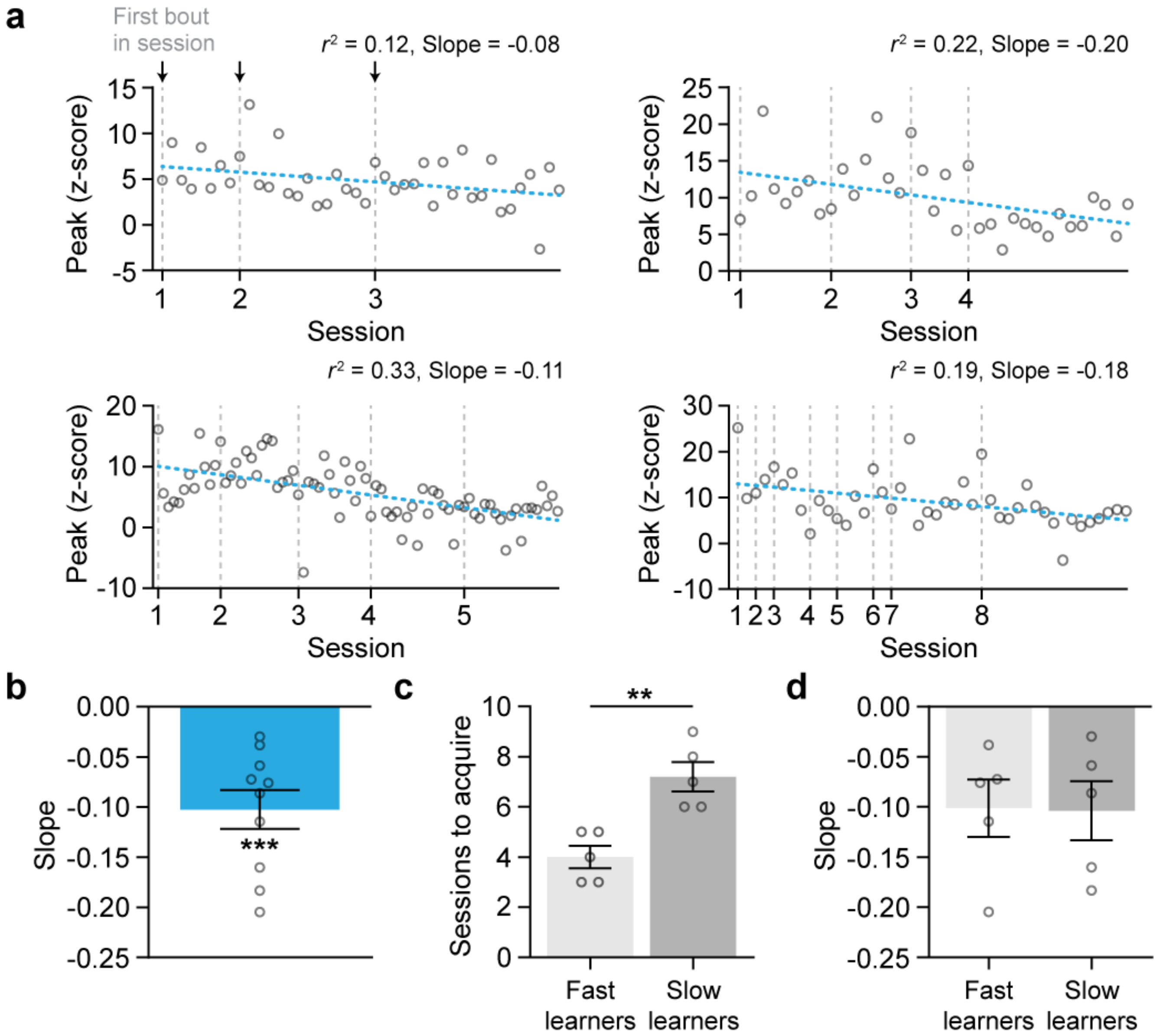


**Extended Data Figure 7. Decrease in mPFC dopamine response during sucrose consumption is independent of learning rate. (a)** Magnitude of mPFC dopamine responses to all lick bouts initiated throughout continuous reinforcement training (4 example subjects represented). Vertical lines indicate the first lick bout in each session. **(b)** Slope values from the linear regression of mPFC dopamine responses to all lick bouts initiated throughout continuous reinforcement training for all 10 subjects tested. The mean slope was negative, indicating that mPFC dopamine responses decreased across sucrose access periods (one sample t-test, $H_0 = 0$, $t_{(9)} = 5.297$, $p < 0.001$). **(c)** Median split demonstrating a differential rate of learning between fast and slow learners (unpaired t-test, $t_{(8)} = 4.355$, $p < 0.01$). **(d)** There was no difference in the slope of the dopamine response during sucrose consumption across access periods between fast and slow learners (unpaired t-test, $t_{(8)} = 0.06018$, $p = 0.9535$). Data represented as mean $\pm$ S.E.M. ** $p < 0.01$; *** $p < 0.001$.

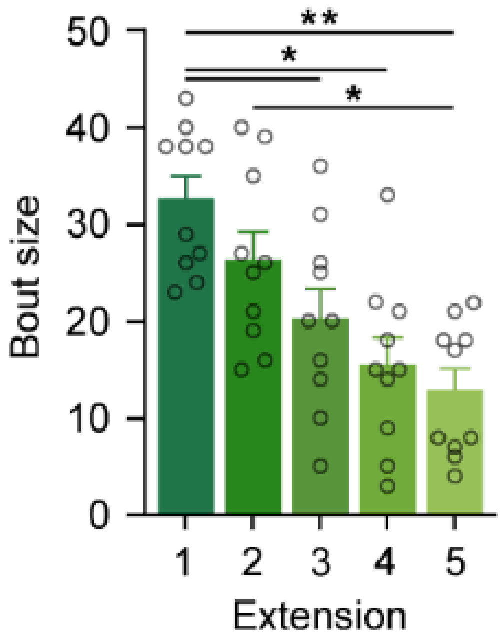


**Extended Data Figure 8. Conditioned licking gradually declines across sipper extensions.** During conditioned reinforcement, animals reliably licked the dry/empty sipper tube during the initial reinforcer presentations, and the size of individual lick bouts decreased across the first 5 sipper extension/reinforcers earned (one-way repeated measures ANOVA, $F_{(2.360,\ 21.24)} = 11.45$, $p = 0.0002$; Tukey's test, 1$^{st}$ vs. 3$^{rd}$, $p = 0.0490$; 1$^{st}$ vs 4$^{th}$, $p = 0.0153$; 1$^{st}$ vs. 5$^{th}$, $p = 0.0052$; 2$^{nd}$ vs. 5$^{th}$, $p = 0.0486$; $p > 0.05$ for all other comparisons). Data represented as mean $\pm$ S.E.M. * $p < 0.05$; ** $p < 0.01$.

a

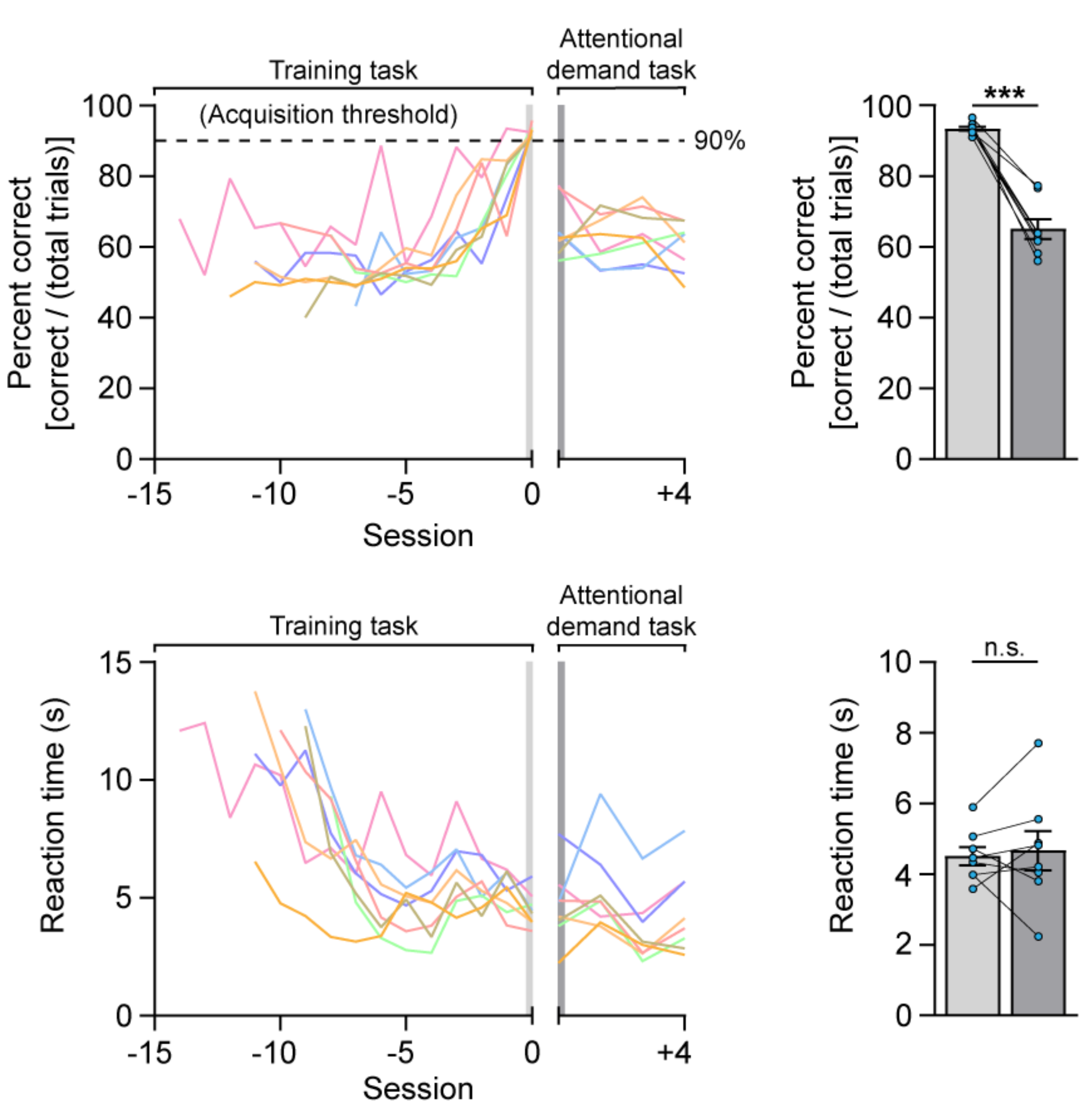


**Extended Data Figure 9. Manipulation of attentional demand affects task performance.** Comparison of behavioral measures during the final session of the training task when animals met acquisition and the first session of the attentional demand task wherein the task parameters were modified to increase attentional demand. Colored lines featured on each line graph represent individual animals. **(a)** Confirming the enhanced difficulty of the attentional demand task, animals achieved a lower percentage of correct trials during the first session of the attentional demand task relative to the final session of the training task (paired samples t-test, $t_{(7)} = 10.76$, $p < 0.001$). **(b)** Reaction time (i.e. time between the extension of the response levers and the response) was not affected by the task switch (paired samples t-test, $t_{(7)} = 0.3813$, $p = 0.7143$). Data represented as mean ± S.E.M. *** $p < 0.001$.
