## Supplemental Video 1 Legend for "Medial prefrontal dopamine dynamics reflect allocation of selective attention"

**Supplemental Video 1. Visualization of mPFC dopamine activity and lick rate throughout sucrose access session.** Replay of an entire 20-minute session of open access to lick spout containing sucrose solution (10% w/v) reveals striking covariance between lick rate (orange) and mPFC dopamine activity (blue). Real time from the start of the session is displayed in the top right of the left panel (note that playback speed is reduced during lick bouts). The fluorescence trace was downsampled from 25 to 6.25 Hz by a 4 times block-wise average, and lick rate was calculated as licks per 160 ms, thus matching the sample rates of the two signals. Both signals were min/max normalized to their respective lowest (0) and highest (1) values throughout the session. No additional processing was performed. The full session is displayed in the left panel and a 30 second sliding window is displayed on the right (the time window displayed on the right is denoted in the full session panel by the grey shading). Pearson’s correlations between lick rate and fluorescence for all animals tested are presented in Extended Data Fig. 4.
